# S-acylation is involved in tonoplast targeting of flax resistance protein M

**DOI:** 10.1101/2020.07.01.181701

**Authors:** Oliver Batistič

**Affiliations:** Institut für Biologie und Biotechnologie der Pflanzen, Universität Münster, Schlossplatz 7, 48149 Münster, Germany, Mail

**Keywords:** R protein M, flax, S-acylation, tonoplast, plasma membrane, PAT10

## Abstract

Several plant resistance proteins require accurate targeting to cellular membranes for effective pathogen defence function. The resistance protein variant M from *Linum ussitatissimum* (flax) protects plants against the flax rust disease and is specifically targeted to the vacuolar membrane. This localization mechanism involves the S-acylation of cysteine residues within the N-terminus of the protein. Moreover, the M S-acylation and targeting signal likely binds to membranes in the absence of lipid modifications and can switch from the vacuolar membrane to the plasma membrane depending on the S-acylation status. Importantly, plasma membrane targeting was observed when the short targeting signal from M was expressed in an *Arabidopsis thaliana* pat10 mutant plant. On the other hand, tonoplast localization of the N-terminal S-acylation domain was reconstituted in the mutant plant upon co-expression of two highly related PAT10 enzymes from flax. In contrast to the Golgi and tonoplast targeted *Arabidopsis* PAT10, the two homologous enzymes from flax mainly localized to the plasma membrane and were partially observed at the vacuolar membrane when expressed in flax as well as in *Arabidopsis* cells. This implicates that S-acylation of flax proteins could occur at the plasma membrane and that the lipid modification is required for subsequent routing to the vacuolar membrane, which could be also involved in the localization mechanism of M.

## Introduction

Plant pathogen invasion and host adaptation is mediated by an intricate interplay between the secreted pathogen effectors and the endogenous disease Resistance (R) proteins of the host. Interestingly, several of these disparate opponents require protein lipid modifications for the full exertions of either virulence or disease control, respectively (Martin *et al*., 2003; Boyle & Martin, 2015). R proteins like Resistance To Pseudomonas Syringae 5 (RPS5) from *Arabidopsis thaliana* or Pseudomonas tomato (Pto) from tomato, as well as various pathogen effectors from *Phytophtora syringae*, utilize N-terminal myristoylation and S-acylation to target to the plant plasma membrane (Boyle & Martin, 2015). N-Myristoylation represents an irreversible modification of an N-terminal glycine residue within a specific consensus sequence (Turnbull & Hemsley, 2017). In contrast to that, S-acylation is the reversible thioesterification of cysteine residues by either palmitate or stearate. This modification often occurs in conjunction with other lipid modifications like N-myristoylation, or in the vicinity of hydrophobic amino acids but lacks a specific sequence requirement (Li & Qi, 2017; Turnbull & Hemsley, 2017). The S-acylation of target proteins is mediated by protein S-acyl transferases (PATs). These multi-pass membrane proteins contain a central aspartate-histidine-histidine-cysteine – cysteine-rich domain (DHHC-CRD), which is required for lipid transfer to target proteins (Politis *et al*., 2005; Rana *et al*., 2018).

In the plant species *Linum usitatissimum* (flax), the five gene loci K, L, M, N and P, confer resistance to the flax rust disease, caused by the basidiomycete fungus *Melampsora lini* (Islam & Mayo, 1990; Lawrence *et al*., 2007). The M resistance locus encodes seven gene variants (named M to M6) which belong to the NOD-like receptor (NLR) protein family (Lawrence *et al*., 2010). The specific gene variant M differs from other locus variations by its unique N-terminal sequence (Lawrence *et al*., 2010). This domain exhibits sequence features which are also present in the N-termini of S-acylated, tonoplast bound Calcineurin B-like (CBL) calcium sensor proteins from *Arabidopsis* (Takemoto *et al*., 2012). It was shown that the first 30 amino acids from M fused to a green fluorescent protein (GFP) confers targeting to the vacuolar membrane but the exact localization mechanism is not fully clarified yet (Takemoto *et al*., 2012). On one hand, direct involvement of S-acylation in the M N-terminus targeting was rather excluded, since the S-acylation inhibitor 2-Bromopalmitate (2-BrP) did not affect the localization of the M N-terminus (Takemoto *et al*., 2012). However, mutation of the cysteine residues within the N terminus resulted in cytosolic accumulation, like it was shown for S-acylated CBLs (Takemoto *et al*., 2012; Batistič *et al*., 2012). Therefore, the potential role of cysteine modification in M targeting can not be completely ruled out.

Here the targeting mechanism and potential S-acylation of flax R protein M by its N-terminal structure were analysed in detail. This work shows that full-length M targeting to the tonoplast and function depends on N-terminal cysteine residues. The first 23 amino acids of M (Mn) represents a minimal S-acylation signal and is sufficient to mediate tonoplast binding in *Nicotiana benthamiana, A. thaliana* and flax cells. The targeting of Mn to the vacuolar membrane can be blocked by the inhibitor 2-BrP, but results in re-targeting of the M N-terminus from the tonoplast to the plasma membrane, indicating that the N-terminal peptide itself could represent a potential membrane-binding domain. Full-length M and Mn can be modified by two highly related flax PAT enzymes homologous to PAT10 from *A. thaliana*, which is known to be involved in the lipid modification of tonoplast *Arabidopsis* CBLs (Zhou *et al*., 2013). Interestingly, in contrast to tonoplast bound *Arabidopsis* PAT10, the flax PAT10 homologues are mainly targeted to the plasma membrane and can retarget Mn-GFP from the plasma membrane to the vacuolar membrane when expressed in the *Arabidopsis* pat10 mutant. This specific localization of the flax PAT10 enzymes could indicate for a specific lipid modification and targeting mechanism of potential substrates at the plasma membrane with subsequent targeting to the vacuolar membrane.

## Material and Methods

### Transient transformation of *Nicotiana benthamiana*, of *Linum ussitatissiumum* and *Arabidopsis thaliana* plants, application of 2-BrP and trypan blue staining, transformation of yeast cells and microscopic analysis

Growth and transient transformation of *N. benthamiana* plants were performed as described previously (Batistič, 2012). For transient *L. ussitatissimum* transformations, seeds were covered with soil and grown for 10-12 days in a growth chamber under a 16h(22°C)/8h(18°C) cycle. Fully expanded cotyledons of *L. ussitatissimum* plants were transformed by direct infiltration of Agrobacteria tumefaciens (GV3101/pMP90). The Agrobacteria suspension was prepared from an overnight grown culture, incubated in activation media (10 mM MES/KOH pH 5.6, 10 mM MgCl_2_ and 600 µM Acetosyringone) for 3 hours (each culture O.D._600_ = 1,5). For microscopic analysis, infiltrated *N. benthamiana* plants were incubated for two days, *L. ussitatissimum* plants for three days.

For the transient transformation of *A. thaliana* seedlings, sterilized seeds were placed on 1x MS media, including MES buffer and 0,7% agar, which was covered with a metal mesh. WT seeds were incubated in a light chamber (Rubarth Apparate GmbH, Laatzen, Germany) at 120-150 µmol/m^2^ /s, under a 16h(22°C)/8h(18°C) day/night regime for four days, pat10 mutants for six days. For transformation, an Agrobacteria suspension was prepared in activation media (see above), mixed with Tween (0,0001 %), and the seedlings were completely submerged in the suspension. The seedlings were incubated for 5 minutes. Subsequently, a vacuum was applied using a water-jet pump (5 minutes) and the vacuum was released (in total three repeats). Afterwards, the seedlings were incubated for further 5 minutes, the suspension was removed and the seedlings were transferred back to the growth chamber for three days.

Application of 2-BrP was performed as described previously using EtOH as the solvent (Batistič *et al*., 2012). Trypan blue staining was performed as described (Nowicki *et al*., 2012). Yeast *Saccharomyces* cerevisiae BY4741 strain (Baker Brachmann *et al*., 1998) were transformed by PEG-heat shock (Albrecht *et al*., 2001) and selected on synthetic dropout media lacking leucine and uracil. To quantify plasma membrane binding in yeast cells, the plasma membrane was stained using CellBrite™ Fix 555 Membrane Stain (Biotium, Freemont, CA, USA). Staining was performed according to the manufacturer’s protocol for at least 10 min at 4°C. Cells were washed two times with fresh media to remove excess stain and subsequently used for microscopic analysis. The given fluorescence ratio is the quotient of the fluorescence intensities at the plasma membrane and within the cell (excluding the plasma membrane and the vacuole). Microscopic analysis of plant and yeast cells were performed using a Leica confocal SP5 system as described (Batistič, 2012).

### Protein S-acylation assays

To determine the S-acylation in plants, proteins were transiently overexpressed in *N. benthamiana* for three days. ∼200 mg of leaf material was frozen in liquid nitrogen and ground using a Mixer Mill. Proteins were extracted in 1x phosphate-buffered saline (PBS) pH 7.4 (Sambrook & Russell, 2000), supplemented with 5 µl/ml protease inhibitor cocktail (PIC) (Sigma Aldrich GmbH, München, Germany), 1 mM EDTA, 1% Triton X-100 and 25 mM n-ethylmaleimide (NEM) for 2 hours at room temperature under continuous rotation. The extract was cleared from the cell debris by centrifugation (2x 5 min, 2000 g, 4°C) and the proteins were precipitated by methanol/chloroform (Wessel & Flügge, 1984). Equal amount of resuspended proteins (in 100 µl 1x PBS pH 7.4, 8 mM Urea, 5 % sodium-dodecyl-sulfate[SDS]) were either incubated with Hydroxylamine (800 mM, pH 7.4) or Tris (45 mM, pH 7.4), containing 1 mM EDTA, 2 µl/ml PIC and 1 mM methoxy-polyethylene glycol (PEG)-maleimide (5 kDa). Proteins were incubated in the solution for one hour under continuous rotation and subsequently precipitated using Methanol/Chloroform. The proteins were resuspended in SDS loading buffer (62,5 mM Tris/HCl pH 6.8, 10% Glycerin, 2% SDS, 1% b-Mercaptoethanol), the protein concentration was determined and equal protein amounts were separated by SDS-PAGE.

To determine S-acylation in yeast cells, yeast were grown overnight in YPD (∼2 ml) in presence of 250 µM 17-Octadecynoic acid (17-ODYA; from a 50 mM stock dissolved in DMSO) at 23°C (controls were supplemented with DMSO). Cells were harvested and washed two times in 1xPBS. For protein extraction, the cell pellet was frozen in liquid nitrogen and ground using a MixerMill. 1xPBS containing 1% Triton X-100, 3µl/ml PIC and 0,03% lithium-dodecyl-sulfate was applied to the ground cell powder and incubated for 10 min at 65°C to extract the proteins. The cell debris was pelleted by centrifugation (10 min, 24000 g, 4°C) and the supernatant was transferred into a new Eppendorf tube. The protein extract was mixed with Tris(benzyltriazolylmethyl)amine (TBTA, final conc. 0,4 mM), ascorbate (final conc. 4 mM), CuSO_4_ (final conc. 4 mM) and Methoxypolyethylene glycol azide (molecular weight 5 kDa, 5-mPEG azide, final conc. 0,6 µg/µl). The mixture was incubated for 45 min at room temperature, and the reaction was stopped by adding 5x SDS-loading dye containing 20 mM EDTA. Equal amounts of proteins were separated by SDS-PAGE.

Blotted proteins were detected using anti-GFP (LifeTechnologies, Darmstadt, Germany) and anti-red FP (RFP) antibody (ChromoTek GmbH, Planegg-Martinsried, Germany) (both, 1:4000 dilution), respectively, and using secondary antibodies coupled with horseradish peroxidase (anti-goat-HRP for anti-GFP, Biorad, München, Germany, 1:10000; anti-rat-HRP for anti-RFP, LifeTechnologies, 1:4000). Detection was performed by chemiluminescence reaction, incubating membranes for 2 minutes in 0,1M Tris-HCl pH 8.6, 1,25mM L-012 (FUJIFILM Wako Chemicals, Neuss, Germany), 0,1 mg/ml para-Hydroxycoumarinsäure (from a 10x stock dissolved in DMSO) and 0,04% H_2_O_2_.

### Phylogenetic analysis

To identify PAT10 homologues, BLASTP searches were performed within the Phytozome database (http://www.phytozome.net/) (Goodstein *et al*., 2012), using the *A. thaliana* PAT10 protein sequence as a query. Multiple protein alignments were performed using ClustalX (Thompson *et al*., 1997). A phylogenetic tree was calculated by a Neighbour Joining method (with 1000 bootstrap replicates) using the MEGA7.0 program (Kumar *et al*., 2008). Pairwise alignment and visualization was performed using BLASTP and Jalview, respectively (Altschul *et al*., 1990; Waterhouse *et al*., 2009).

### cDNA preparation from *Linum ussitatissimum* seedlings and construct generation

For cDNA preparation, flax seeds were sown in soil and seedlings were grown in the dark for 7 days at 21°C. RNA was prepared from etiolated cotyledons and hypocotyl (∼300 mg), using the SV Total RNA Isolation System (Promega GmbH, Mannheim, Germany). For cDNA synthesis, 1 µg RNA was used in a reverse transcription reaction as described previously (Batistič, 2012).

The sequence of all primers to generate the used constructs are given in Table S1. A detailed description of the constructs generated is given in the Supplemental Material and Method.

### Accession numbers

LuM (GenBank: U73916.1), LuPAT10a (Phytozome: Lus10028384), LuPAT10b (Phytozome: Lus10041837), avrM (GenBank: ABB96264), AvrM-A (GenBank: ABB96258), AtPAT10 (At3g51390), AtPAT11 (At3g18620), AtTPK1 (At5g55630), AtCBL2 (At5g55990), AtCBL3 (At4g26570), AtCBL6 (At4g16350)

## Results

### M function and tonoplast targeting requires intact N-terminal cysteines and M localization is affected by 2-BrP

Several NLR proteins require membrane binding for correct function, which could be mediated by protein lipid modifications (Gao *et al*., 2011; Qi *et al*., 2012; Takemoto *et al*., 2012). It was shown that the N-terminus of M mediates tonoplast binding and that this depends on cysteine residues (Takemoto *et al*., 2012). Whether the N-terminus also mediates the vacuolar membrane targeting of the full-length M protein, and which mechanisms underlie this localization are not fully understood. Therefore, the localization of full-length M protein fused to GFP (M-GFP) was analysed by transient expression in *N. benthamiana* plants and compared to the plasma membrane marker protein TM23 (fused with RFP, TM23-RFP) (Batistič *et al*., 2008). To facilitate detectable levels of M-GFP, the expression was driven from an Estradiol inducible vector system (Schlücking *et al*., 2013). In leaf samples without Estradiol, only the red fluorescence from the TM23-RFP fusion was detectable at the plasma membrane, while green fluorescence was observed only in samples incubated with Estradiol (Fig. **1a**). However, this fluorescence of M-GFP was observed at the internal membrane which was separated from TM23-RFP (Fig. **1a**), indicating that M is targeted to the vacuolar membrane. To confirm this observation, M-GFP was co-transformed with the vacuolar marker TPK1-RFP (Batistič, 2012). In presence of Estradiol, M-GFP fluorescence was observed overlapping with TPK1-RFP (Fig. **1b**), corroborating that M is targeted to the tonoplast likely mediated by the N-terminus (Takemoto *et al*., 2012). It was previously shown that mutations of the cysteines 19 and 20 (and of glycine 21) affect the membrane binding of the M N-terminus (Takemoto *et al*., 2012). Therefore, to investigate if other protein domains of M are involved in membrane binding, the N-terminal cysteines were mutated to alanine. This protein was fused to GFP (M^C19,20A^-GFP) and expressed in *N. benthamiana* induced by Estradiol. This revealed that the mutated protein accumulated in the cytosol and nucleus (Fig. **1c**), showing that the full-length M requires intact N-terminal cysteine residues for efficient membrane binding which is in accordance to the results using the isolated N-terminus (Takemoto *et al*., 2012). This approved that the targeting of M is rather mediated by the N-terminus and that other domains of M are unlikely to be involved or play a minor role in the localization mechanism. To determine if S-acylation might be involved in the targeting of M, full-length M was expressed in the presence of the S-acylation inhibitor 2-BrP. Surprisingly, M was not targeted to the vacuolar membrane but accumulated in the cytosol and nucleoplasm like the mutated M^C19,20A^ protein (Fig. **1d**), while TM23-RFP localization to the plasma membrane was not affected (Fig. **1d** and S1).

**Figure 1:**
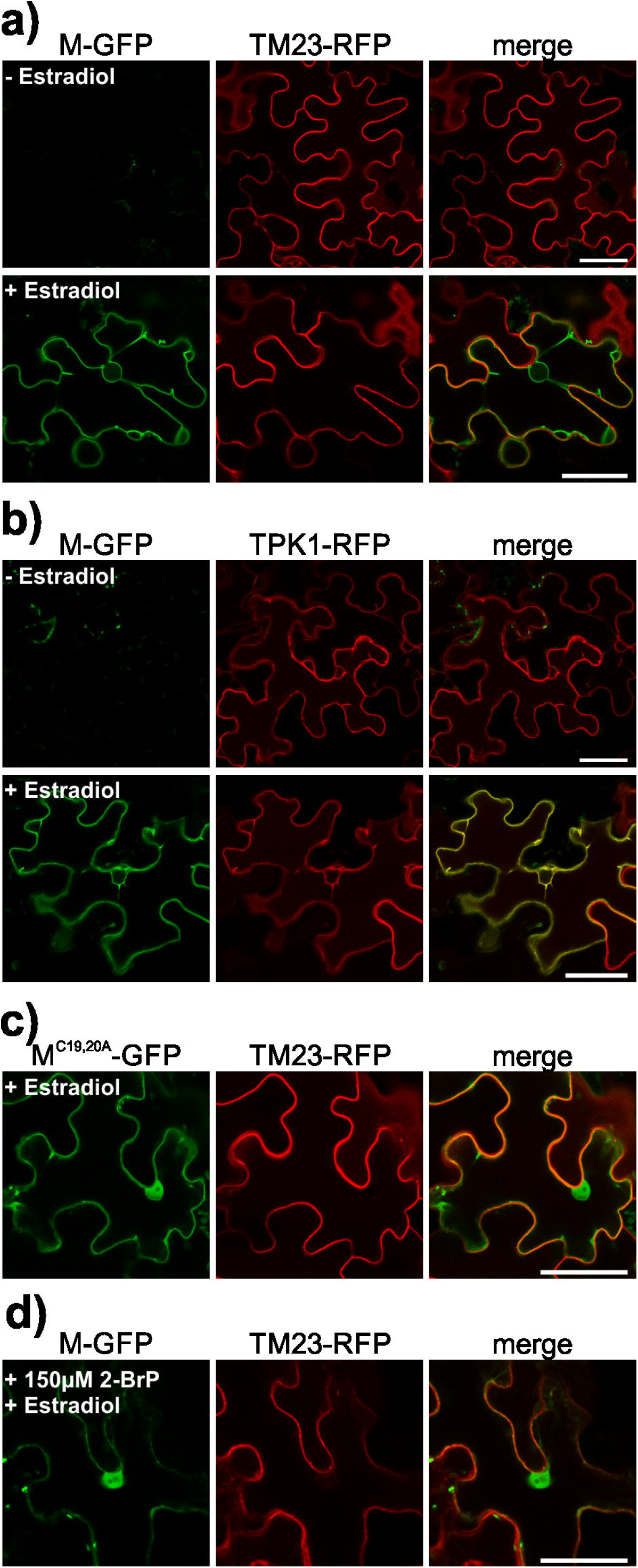
Tonoplast binding of full-length M requires N-terminal cysteines and is affected by 2-BrP. Full-length M and the M^C19,20A^ mutant, fused to GFP (M-GFP and M^C19,20A^-GFP) (GFP is shown in the first image), were expressed in *N. benthamiana* leaves. Expression was induced by Estradiol application (125 µM Estradiol, containing 0,1% EtOH) three days after Agrobacteria infiltration and microscopic analysis were performed 24 h after Estradiol application (pictures shown in the second row in **a** and **b**; a sample lacking Estradiol is shown in the first row as a control). The plasma membrane marker TM23-RFP or the vacuolar marker TPK1-RFP (second image in **a** and **b**, respectively) were co-expressed using a constitutive promoter. Full-length M-GFP is targeted to the vacuolar membrane while TM23-RFP is localized to the plasma membrane (both fluorescences merged in the third image) (**a**). In contrast, M-GFP co-localized with the vacuolar marker TPK1-RFP (**b**) and M^C19,20A^-GFP accumulates in the cytosol (**c**). Similarly to that, the application of 150 µM 2-BrP results also in cytoplasmic accumulation of M-GFP (**d**). The scale bars depicted represent 50 µm.

To further test if the presence of N-terminal cysteines in M is also required for inducing a hypersensitivity response (HR), M and M^C19,20A^ were expressed either in presence of the active effector AvrM-A or the non-active effector avrM from *M. lini*, both lacking the N-terminal signal peptide (ΔSP) (Catanzariti *et al*., 2006). As shown, after application of Estradiol to induce expression of M or M^C19,20A^, only M in combination with AvrM-AΔSP induced an HR in the infiltrated region, while co-expression of M^C19,20A^ together with AvrM-AΔSP, or the expression of M/M^C19,20A^ with avrMΔSP, did not induce an HR (Fig. **2**).

**Figure 2:**
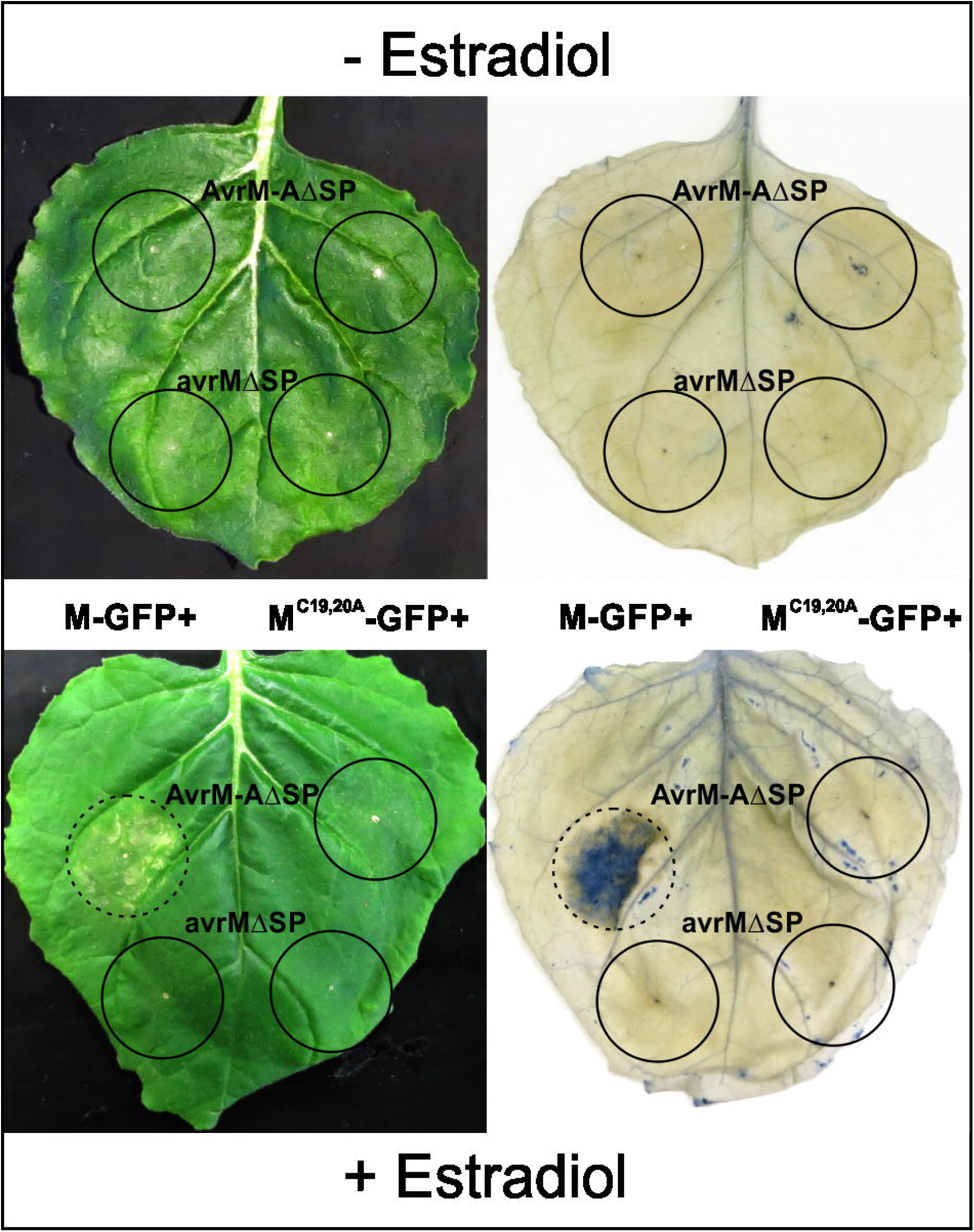
N-terminal cysteines in full-length M are required to induce cell death in combination with pathogen effectors. Full-length M (left half of the leaf) and the M^C19,20A^ mutant (right half of the leaf) (both fused to GFP: M-GFP and M^C19,20A^-GFP), were co-expressed with the AvrM-AΔSP (upper half) or with the avrM-ΔSP (lower half) effector proteins from *M. lini* (expressed using a constitutive promoter). The upper leaf was infiltrated with the respective combinations of Agrobacteria, but Estradiol was omitted. In the lower leaf, Estradiol was applied 3 days after infiltration to induce M-GFP/M^C19,20A^-GFP expression, and cell death was recorded after 7 days. Only the combination of full-length M together with AvrM-AΔSP induced cell death after Estradiol application (left image and right image, indicated by dashed circles), as indicated by the trypan blue staining (right image).

Taken together, these results show that M targets to the vacuolar membrane and that this requires intact N-terminal cysteines, approving that the N-terminus represents an important targeting signal. The presence of the N-terminal cysteines is also a prerequisite to induce an HR by M in the presence of an effector, indicating that tonoplast targeting is required for M function. Importantly, the targeting of M was affected by the inhibitor 2-BrP and results in cytoplasmic accumulation of the protein. This is similar to results known from S-acylated, tonoplast bound CBLs from *Arabidopsis* which accumulate in the cytosol in presence of 2-BrP as well (Batistič *et al*., 2012; Zhang *et al*., 2017). This, therefore, indicates that N-terminal S-acylation is likely involved in M targeting and function at the tonoplast.

### The first 23 amino acids from M are sufficient for tonoplast binding and S-acylation but relocates to the plasma membrane in the presence of 2-BrP

To determine if M targeting directly involves S-acylation of the N-terminus, the localization of the M N-terminal (Mn) domain in absence and presence of the inhibitor 2-BrP was re-assessed. The flax M and the *Arabidopsis* CBL2/3/6 N-termini exhibit several overlapping structural features (Fig. **3a**) (Takemoto *et al*., 2012). Especially the two N-terminal cysteines residues of M and a central hydrophobic stretch are similarly arranged in Mn and CBL6n, indicating that the cysteines of M could represent targets for lipid modification. This presumption is further strengthened using the S-acylation prediction programme CSS-PALM (Ren *et al*., 2008) which revealed a high probability of potential lipid modification (probability score: 10,56 and 18,89 for cysteine 19 and 20, respectively). To determine if targeting of the M N-terminus is affected by 2-BrP, the first 23 amino acids of Mn, which correspond to the minimal targeting signal in CBL2/CBL3 (Batistič *et al*., 2010), were N-terminally fused to GFP (Mn-GFP). The localization of Mn-GFP was compared to the tonoplast protein TPK1-RFP which is unaffected by 2-BrP (Batistič *et al*., 2012). In the EtOH control solution, Mn-GFP targeted to the vacuolar membrane and co-localized with TPK1-RFP in *N. benthamiana* cells (Fig. **3b**). However, in samples incubated with 150 µM 2-BrP, TPK1 associated with the vacuolar membrane, but Mn-GFP did not accumulate in the cytosol like the full-length M protein. Surprisingly, Mn-GFP retargeted from the tonoplast to the plasma membrane while a minor fraction remained at inner membrane structures. To further confirm plasma membrane re-targeting, Mn-GFP was co-expressed with the plasma membrane marker TM23-RFP in presence of EtOH or 150 µM 2-BrP. In the solvent control, Mn-GFP and TM23-RFP displayed differential localization at the vacuolar and the plasma membrane, respectively. In contrast, in the presence of 2-BrP Mn-GFP strongly co-localized with the plasma membrane marker (Fig. **3c**), indicating that S-acylation is involved in the correct targeting of the M N-terminus. To further determine the role of the cysteines in membrane binding and/or targeting of Mn, the potential S-acylation sites (cysteines 19 and 20) were mutated to alanine and fused with GFP (Mn^C19,20A^-GFP). This mutated protein accumulated in the cytosol/nucleoplasma and did not displayed binding to membranes showing that the cysteines are required for Mn membrane binding in plant cells (Fig. **3d**).

**Figure 3:**
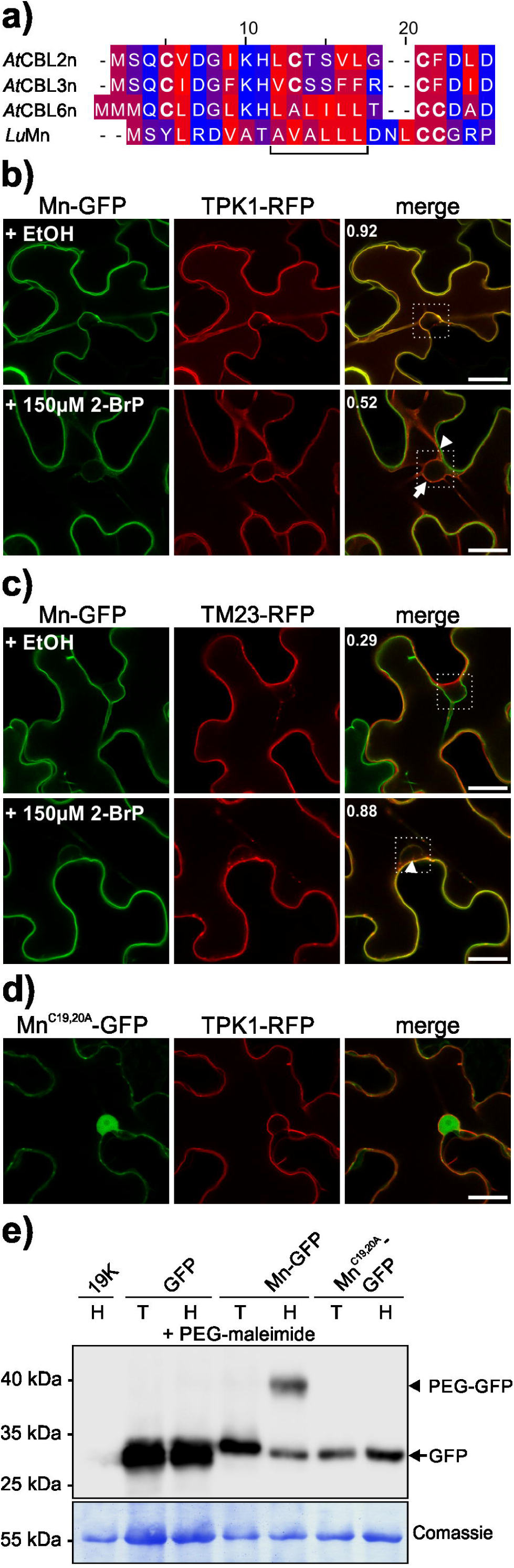
The M N-terminus relocates from the tonoplast to the plasma membrane by 2-BrP, and intact cysteines are required for membrane binding and S-acylation. **a)** Amino acid sequence alignment of the N-termini of CBL2, CBL3, CBL6 from *Arabidopsis thaliana* (*At*CBL2/3/6-n) and R protein M from *Linum ussitatissimum* (*Lu*Mn). The colouring displays the hydrophobicity of the amino acids (hydrophilic blue, lipophilic red). The horizontal bracket indicates a hydrophobic stretch in AtCBL6-n and *Lu*Mn. Cysteines are highlighted in bold letters. **b)** M N-terminus (first 23 amino acids), fused to GFP (Mn-GFP) (first column), was co-expressed with the tonoplast marker TPK1 fused to RFP (TPK1-RFP) (second column) in *N. benthamiana* leaves in presence of EtOH control solution (first row), or 150 µM S-acylation inhibitor 2-BrP (second and third rows). In presence of the EtOH control solution, both fluorescences overlap at the vacuolar membrane (merged image in the third column). In the presence of 2-BrP, Mn-GFP is shifted to the plasma membrane (arrowhead) while TPK1-RFP localizes to the tonoplast (arrow). The co-localization of both fluorescences was determined within the marked box (indicated in the merged image), the number indicates the Pearson’s correlation coefficient. The scale bars depicted in the overlay images are 20 µm. **c)** Mn-GFP (first column) was co-expressed with the plasma membrane marker TM23 fused to RFP (TM23-RFP) (second column) in *N.benthamiana* leaves, in presence of EtOH control solution (first row), or 150 µM 2-BrP (second row). In the presence of the EtOH control solution, both fluorescences show weak overlap (merged image in the third column). In the presence of 2-BrP Mn-GFP is shifted to the plasma membrane (arrowhead) indicated by the increased co-localization with TM23-RFP. The co-localization of both fluorescences was determined within the marked box (indicated in the merged image), the number indicates the Pearson’s correlation coefficient. The scale bars in the overlay images represent 20 µm. **d)** Microscopic analysis of the mutated Mn-GFP, where both N-terminal cysteine residues were replaced by alanine (Mn^C19,20A^-GFP, first image), co-expressed with TPK1-RFP (second image) in *N. benthamiana* leaves (merged in the third image). Mn^C19,20A^-GFP exhibits cytoplasmic localization while TPK1 is targeted to the vacuolar membrane. The scale bar in the overlay image is 20 µm. **e)** S-acylation of Mn-GFP determined by an S-acyl-PEG switch assay. GFP as a control, Mn-GFP, and Mn^C19,20A^-GFP were expressed in *N. benthamiana* leaves. Extracted proteins were incubated in NEM subsequently either in Tris (control solution, T) or Hydroxylamine solution (H) in the presence of PEG-maleimide. A leaf expressing only the helper plasmid 19K (incubated with Hydroxylamine) was used as an antibody specificity control. GFP proteins were detected in all samples (except the 19K control) at the expected size (27-30 kDa, free GFP and Mn-GFP, respectively; arrow) (upper part). The Mn-GFP sample incubated with Hydroxylamine exhibits a further, shifted band at ∼40 kDa (arrowhead), indicating S-acylation of the M N-terminus. Coomassie staining of the rbcL protein (lower blot) is shown as a loading control. The position of a protein size marker is indicated.

These results showed that the N-terminus of M represents a potential lipid modification signal, which when mediates the correct targeting of M. Therefore, the S-acylation of the M N-terminus was assessed biochemically, using the S-Acyl-PEG switch assay (Percher *et al*., 2016). In this assay, free thiol groups of a protein are first blocked using NEM, followed by removal of cysteine-thioester groups using Hydroxylamine. The now free thiol groups are subsequently linked with PEG-maleimide, which results in a protein mass increase. To determine the S-acylation of the M N-terminus, the 19K protein (immunoblot control), unmodified GFP, Mn-GFP and Mn^C19,20A^-GFP were transiently expressed. The proteins were extracted in the presence of NEM and afterwards incubated either in Tris control solution or Hydroxylamine both containing PEG-maleimide. After SDS-PAGE separation, a GFP was detected in all samples (except in the 19K Hydroxylamine control sample) at the expected size (27-30 kDa, GFP and Mn-GFP, respectively) on the immunoblot (Fig. **3e**). Importantly, only Mn-GFP incubated in the Hydroxylamine solution, displayed a further shifted protein (∼40 kDa), which was not observed in the Tris control sample, showing that Mn was S-acylated in *N. benthamiana* cells. The GFP control, as well as the mutant Mn^C19,20A^-GFP, neither displayed a shifted protein in the Tris control nor the Hydroxylamine solution (Fig. **3e**), showing that these proteins were not S-acylated and therefore not modified by PEG-maleimide.

These results proved that the first 23 amino acids of M are sufficient for tonoplast targeting and S-acylation. However, in the presence of 2-BrP, this specific Mn peptide retargeted from the vacuolar membrane to the plasma membrane. This finding is in contrast to a previous result, there no effect of 2-BrP was observed when using the first 30 amino acids of M (Takemoto *et al*., 2012). These differences could rely on an additional function of the amino acids 24-30 in the membrane binding and targeting efficiency of the M N-terminus. To test this, the first 30 amino acids of M were fused to GFP (Mn_30_-GFP) and the effect of 2-BrP was tested using the same concentration as previously reported (100 µM 2-BrP) (Takemoto *et al*., 2012). In the control solution, Mn_30_-GFP targeted to the tonoplast and did not co-localize with the plasma membrane marker TM23-RFP. On the other hand, in the presence of 100 µM 2-BrP, a partial fraction of Mn_30_-GFP localized to the plasma membrane but in contrast to the shorter 23 amino acid fragment of M, a substantial fraction of Mn_30_-GFP remained at the vacuolar membrane (Fig. S2A). This indicates that amino acids 24-30 either promote vacuolar membrane binding or somehow reduced the effect of 2-BrP. Consequently, only a minor fraction of the longer M N-terminus targets to the plasma membrane in the presence of 2-BrP. Therefore, to affirm that Mn_30_-GFP is lipid-modified as well, the S-acylation of Mn_30_-GFP was tested using the PEG-switch method. This revealed that Mn_30_-GFP shifted in the presence of Hydroxylamine and PEG-maleimide but not in the Tris control, similar to the shorter Mn fragment (Fig. S2B). This result finally corroborated that the Mn_30_ peptide is S-acylated and partially shifted to the plasma membrane in presence of 2-BrP.

Altogether, these analyses showed that the N-terminus of M is sufficient to mediate vacuolar membrane binding. Biochemical analysis on two variants of the M N-terminus revealed that this domain is S-acylated in plant cells. Surprisingly, in the presence of the S-acylation inhibitor 2-BrP, Mn-GFP and Mn_30_-GFP did not accumulate in the cytosol like the full-length M, and especially Mn-GFP retargeted from the vacuolar membrane to the plasma membrane. This indicates that the M N-terminus represents a potential membrane-binding domain and that targeting of the domain could be modulated by S-acylation.

### Lipid modification of Cysteine 19 is sufficient to mediate vacuolar membrane binding of Mn

To further determine if either one cysteine or both N-terminal cysteines are involved in the lipid modification and targeting of M, single cysteine to alanine mutations were introduced into Mn-GFP (cysteine 19 or 20 to alanine) and the localization of the peptides in absence and presence of 2-BrP were tested. (Fig. **4**). Interestingly, Mn ^C19A^ -GFP and Mn ^C20A^ -GFP showed differential targeting mechanisms, also compared to the intact Mn peptide. Mn ^C19A^ localized to the cytosol in the presence of EtOH and 2-BrP but partially accumulated on moving vesicles (Fig. **4a**). In contrast, Mn ^C20A^ targeted to the vacuolar membrane in the control solution, but in the presence of 2-BrP accumulated in the cytosol and to moving vesicles, similar to Mn ^C19A^ (Fig. **4b**). A PEG-switch was then applied to determine how single cysteine mutations affect S-acylation. Interestingly, both Mn ^C19A^ and Mn ^C20A^ displayed a shifted protein in presence of Hydroxylamine and PEG-maleimide like Mn, showing that all proteins could be S-acylated (Fig. **4c**). However, the intensity of the shifted protein is different in the three samples (Mn>Mn ^C20A^ >Mn ^C19A^). This decreasing intensity of the shifted protein in Mn ^C20A^ and Mn ^C19A^ compared to Mn indicates that the efficiency of S-acylation is affected if one cysteine is mutated. However, while Mn ^C20A^ still can bind to the tonoplast, the weak modification of Mn ^C19A^ was not sufficient to detect membrane binding in microscopic analysis. This indicates that cysteine 20 plays a minor role in the targeting mechanism but likely promotes the reactivity of cysteine 19, which seems to be more important in the efficient tonoplast localization of Mn.

**Figure 4:**
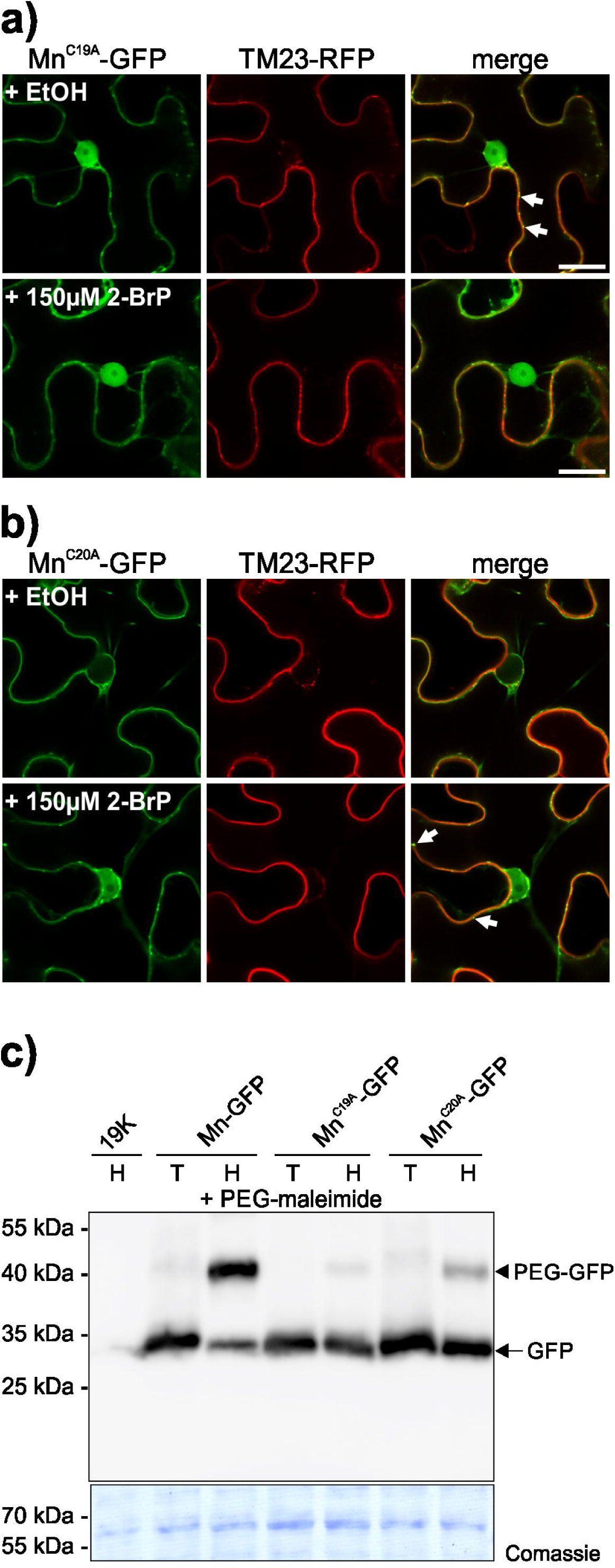
Lipid modification of cysteine 19 can be sufficient to mediated tonoplast targeting of Mn. **a)** M N-terminus containing a cysteine 19 to alanine mutation (fused to GFP: Mn^C19A^-GFP; first column), was co-expressed with TM23-RFP (second column) in *N. benthamiana* leaves in presence of EtOH control solution (first row), or 150 µM 2-BrP (second row). In presence of the EtOH or 2-BrP, Mn^C19A^-GFP accumulated in the cytosol and partially to vesicles (arrows), while TM23 associated with the plasma membrane (merged image in the third column). The scale bars in the overlay images represent 20 µm. **b)** M N-terminus containing a cysteine 20 to alanine mutation (fused to GFP: Mn^C20A^-GFP; first column), was co-expressed with TM23-RFP (second column) in *N. benthamiana* leaves in presence of EtOH control solution (first row), or 150 µM S-acylation inhibitor 2-BrP (second row). In presence of the EtOH control solution, Mn^C20A^-GFP was associated with the vacuolar membrane while TM23 associated with the plasma membrane (merged image in the third column). In presence of 2-BrP Mn^C20A^-GFP accumulated in the cytosol and partially to vesicles (arrows), but did not associate with the plasma membrane. The scale bars in the overlay images represent 20 µm. **c)** S-acylation of Mn-GFP, Mn^C19A^-GFP and Mn^C20A^-GFP determined by an S-acyl-PEG switch assay. A leaf expressing only the helper plasmid 19K (incubated with Hydroxylamine) was used as an antibody specificity control. GFP proteins were detected in all samples (except the 19K control) at the expected size (∼ 30 kDa) (upper part). In all samples expressing the GFP fusion, a shifted band is observed only in Hydroxylamine sample (H) at ∼40 kDa (arrowhead) but not in the Tris control (T), indicating S-acylation of the fusion proteins. Coomassie staining of the rbcL protein (lower blot) is shown as a loading control. The position of a protein size marker is indicated.

### Two flax PAT10 like proteins can mediate S-acylation of Mn and membrane targeting of full-length M in yeast cells

The obtained results showed that the flax M N-terminus is S-acylated in plant cells and that the lipid modification is required for correct targeting to the vacuolar membrane. In *A. thaliana*, the tonoplast/Golgi localized PAT10 is involved in the targeting of CBL2/3/6, which accumulate in the cytosol when overexpressed in a pat10 mutant (Zhou *et al*., 2013). The targeting and sequence similarities between M and CBL2/3/6 N-termini (Fig. **3**) indicated that a PAT10 homologous enzyme from flax could be involved in the M lipid modification event.

To identify potential PAT10 proteins in flax, the *A. thaliana* PAT10 protein sequence (At3g51390) was used as a query for BLAST searches within the *L. ussitatissumum* proteome. The obtained protein sequences were then used for further alignment and phylogenetic analysis. By this, two sequence annotations (Lus10028384 and Lus10041837) clustered together with *Arabidopsis* PAT10, while the remaining sequences grouped with the other *Arabidopsis* PATs (Fig. S3A). These two sequences are highly related (97% similarity within the protein-coding DNA region) and the proteins differ in only nine amino acid positions (Fig. S3B), indicating that these genes arose by a rather recent duplication event. Both proteins comprise 346 amino acids (∼39,6 kDa), which share 57/58% identical and 73/74% similar amino acids compared to the *Arabidopsis* PAT10 protein (Fig. S3B, S3C). cDNA clones were isolated for both annotated sequences showing that the two potential PAT10 enzymes are expressed in flax, which were subsequently designated as LuPAT10a (Lus10028384) and LuPAT10b (Lus10041837), respectively.

It was recently shown that CBL2 is targeted to the yeast cytosol, implicating that this protein is not efficiently modified by the yeast PAT machinery (Zhang *et al*., 2017). This led to the assumption that yeast cells could be used as a tool to directly test the lipid modification of one target by one plant enzyme *in vivo*. Therefore, to test if yeast cells are suitable to investigate the lipid modification of Mn-GFP, a direct S-acylation assay was conducted (Fig. **5a**). For this, yeast cells were grown in the presence of 17-ODYA, a fatty acid alkyne which can be integrated into proteins by S-acylation (Martin & Cravatt, 2009). To finally detect the thioesterified 17-ODYA-protein, the extracted proteins are modified in a copper-mediated “click-reaction” with 5-mPEG azide, which specifically reacts with the alkyne group of the incorporated 17-ODYA. This then results in a mass shift of the analysed protein, which is detected by SDS-PAGE analysis, similarly to the PEG-switch method described above. When yeast cells expressing Mn-GFP were grown in media supplemented with 17-ODYA, a protein of the expected size (∼ 30 kDa) was identified while a specifically shifted band was undetectable (Fig. **5a**, second lane). This indicated that yeast PAT enzymes were not efficiently modifying the M N-terminus. Furthermore, Mn-GFP appeared as a 30 kDa protein in yeast cells which co-expressed LuPAT10a (third lane) or LuPAT10b (lane six) in the absence of 17-ODYA. In contrast to that, a specifically shifted band (between 40-55 kDa) appeared in yeast samples co-expressing LuPAT10a (lane four) or LuPAT10b (lane seven) in presence of 17-ODYA, showing that here both PATs were able to incorporate 17-ODYA into Mn-GFP, which was then modified by 5-mPEG azide in the “click reaction”. Importantly, co-expression of non-active LuPAT10a/b proteins with a cysteine to alanine mutation in the central DHHC site (LuPAT10a/b^DHHA^) did not affect Mn-GFP size, which appeared as a single 30 kDa protein (lane five and eight). Furthermore, the non-S-acylatable Mn^C19,20A^-GFP fusion was not modified by 17-ODYA and appeared as a single protein as well, although LuPAT10a/b were co-expressed (lane nine and ten). This shows that active LuPAT10a or LuPAT10b can mediate lipid modification of Mn-GFP and that the modification depends on the N-terminal cysteines.

**Figure 5:**
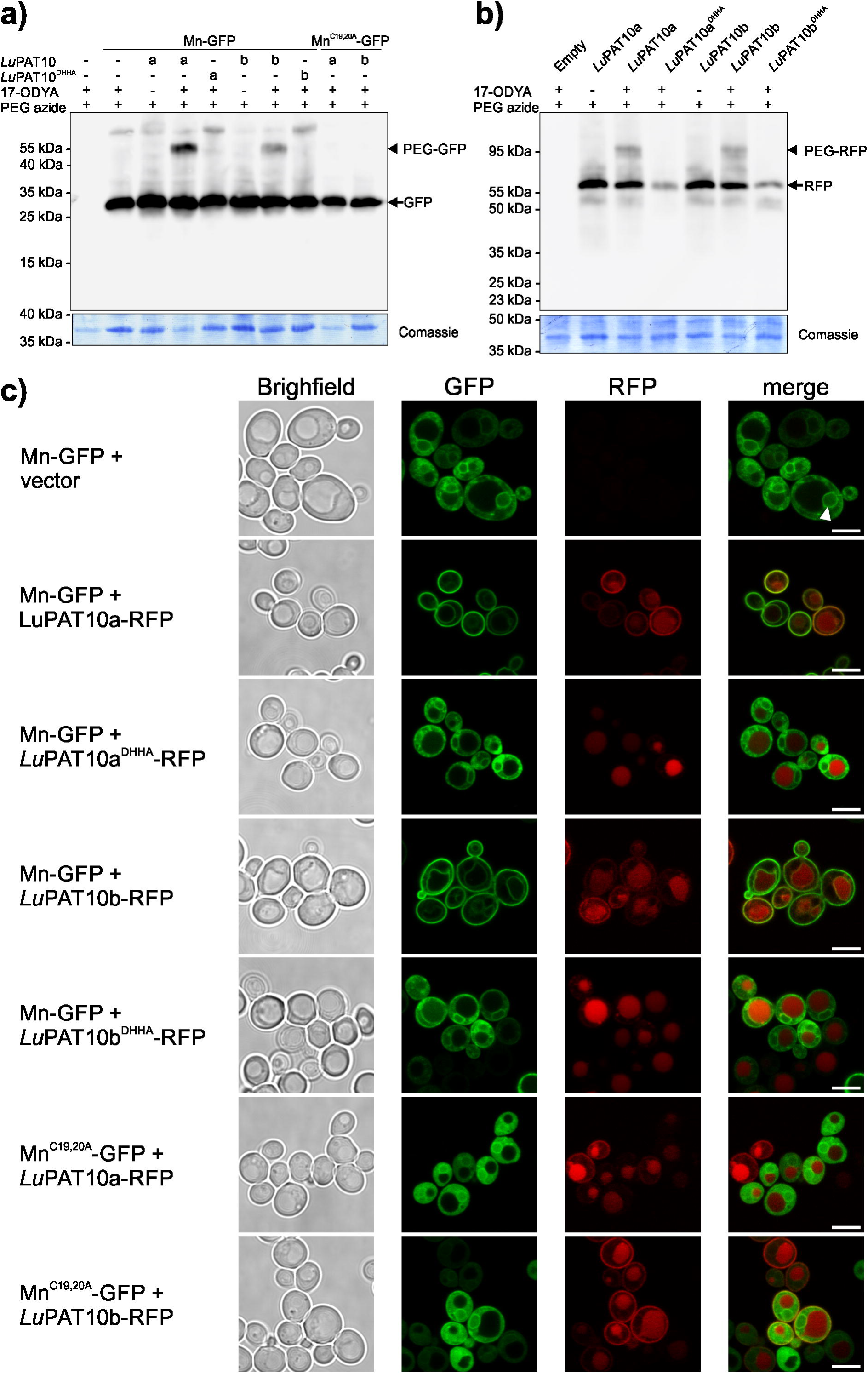
The M N-terminus is S-acylated in yeast cells co-expressing flax PAT10. **a)** S-acylation of Mn-GFP by LuPAT10a/b was determined in yeast cells by analysing the incorporation of 17-ODYA into Mn. Yeast cells were transformed with two empty plasmids (pVT-U + pGVac8Leu; immunoblot specificity control, first lane), with Mn-GFP (in pVT-U) co-transformed with empty vector (pGVac8Leu, second lane), Mn-GFP with LuPAT10a-RFP (in pGVac8Leu, lane three-four), Mn-GFP with LuPAT10a ^DHHA^-RFP (in pGVac8Leu, lane 5), Mn-GFP with LuPAT10b-RFP (in pGVac8Leu, lane six-seven), Mn-GFP with LuPAT10b ^DHHA^-RFP (in pGVac8Leu, lane eight), Mn^C19,20A^-GFP with LuPAT10a-RFP (lane nine) or LuPAT10b-RFP (lane ten). Proteins were extracted and incubated with 5-mPEG azide (PEG azide). Mn-GFP or Mn^C19,20A^-GFP was detected in all samples (except specificity control, first lane) at the expected size (∼30 kDa, GFP; arrow). A specifically shifted protein of Mn-GFP (position indicated by arrowhead) is detectable when either LuPAT10a (lane four) or LuPAT10b (lane seven) were co-transformed and only if 17-ODYA was present. Controls without 17-ODYA (lane three and six) lack the shifted protein. In contrast, Mn^C19,20A^-GFP is not shifted in the presence of 17-ODYA, although PAT10a or PAT10b were co-transformed (lane nine and ten). Coomassie staining of blotted proteins is shown below as loading control. The position of a protein size marker is indicated. b) S-acylation of LuPAT10a and LuPAT10b were determined in yeast cells by analysing the incorporation of 17-ODYA into the enzymes. Yeast cells transformed with two empty plasmids (immunoblot specificity control, first lane), expressing LuPAT10a (lane two-three), the non-active LuPAT10a^DHHA^ (lane four), LuPAT10b (lane five-six) or the non-active LuPAT10b^DHHA^ (lane eight), were grown in presence of 17-ODYA (controls lacking 17-ODYA are shown in lane two and five). Proteins were extracted and incubated with 5-mPEG azide (PEG azide). A PAT-RFP fusion is detectable in all samples (∼ 60 kDa, arrow) except in the vector control (first lane). In presence of 17-ODYA, a shifted protein is detectable for LuPAT10a (lane three) and LuPAT10b (lane six) (∼ 95 kDa, indicated by the arrowhead). This shifted protein is not detectable when either LuPAT10a^DHHA^ (lane four) or the non-active LuPAT10b^DHHA^ (lane eight) were expressed. Coomassie staining of blotted proteins is shown below as loading control. The position of a protein size marker is indicated. **c)** Microscopic analysis of yeast cells expressing Mn-GFP. In yeast cells, Mn-GFP is targeted partially to the ER (arrowhead) and the cytosol. Co-expression of LuPAT10a-RFP (row two) or LuPAT10b-RFP (row four) (both mainly detected within the yeast vacuole, partially at the plasma membrane) results in targeting of Mn-GFP to the yeast plasma membrane and partially to the yeast tonoplast. In contrast, co-expression of mutated LuPAT10a^DHHA^-RFP (row three) or LuPAT10b^DHHA^-RFP (row five) (both mainly detected within the yeast vacuole) does not affect the localization of Mn-GFP. Also, mutated Mn^C19,20A^-GFP partially targets to the yeast nuclear envelope and strongly accumulates in the cytosol, even when LuPAT10a-RFP or LuPAT10b (lane six and seven) are co-expressed. Brightfield pictures are shown in the first column, GFP is shown in the second, RFP in the third, a merged picture of the fluorescences in the last column. Bar represents 5 µm.

The expression and S-acylation of LuPAT10a/b and LuPAT10a/b^DHHA^ in yeast cells were also tested using the direct S-acylation assay (Fig. **5b**). Both, LuPAT10a and LuPAT10b (lane two and five), appeared as a ∼60 kDa RFP fusion protein when expressed in yeast cells lacking 17-ODYA, as expected. In contrast to that, a shifted LuPAT10a/b protein (∼ 95 kDa) was detected in yeast samples, which were grown in 17-ODYA supplemented media (lane three and lane six). On the other hand, this shifted 95kDa protein was not detectable when the inactive LuPAT10a/b ^DHHA^ proteins were expressed in presence of 17-ODYA, showing that these proteins were not S-acylated efficiently (neither in cis nor in trans). Moreover, the weaker bands of LuPAT10a/b^DHHA^ further indicated that the proteins were either less well expressed or less stable than the active wild-type proteins, further showing that the activity status could determine the protein level of these enzymes.

These yeast cells were then used for microscopic analysis. Mn-GFP expressed in yeast cells mainly showed cytoplasmic fluorescence and accumulations around the nucleus (Fig. **5c**, first row). In contrast to that, in cells co-expressing LuPAT10a-RFP (second row) or LuPAT10b-RFP (fourth row), Mn-GFP targeted to the yeast plasma membrane and partially to the vacuolar membrane. This localization of Mn-GFP was not observed when LuPAT10a/b^DHHA^-RFP was co-expressed (row three and five), showing that the plasma membrane targeting of the M N-terminus is mediated by active LuPAT10 enzymes. Furthermore, the mutant Mn^C19,20A^-GFP, which was co-expressed with active LuPAT10a/b-RFP (row six and seven), accumulated in the cytosol and partially around the nucleus similar to Mn. Notably, the fluorescence of LuPAT10a/b and of the mutant versions LuPAT10a/b ^DHHA^ was observed mainly within the vacuoles (Fig. **5c**), implicating that both proteins are targeted for degradation into the yeast vacuole. However, in contrast to the inactive mutants, some fluorescence of active LuPAT10a/b-RFP was also observed at the plasma membrane, indicating that the activity of the enzymes could regulate the targeting efficiency of the enzymes to the yeast plasma membrane.

To further determine the specificity of this modification, Mn-GFP was co-transformed together with LuPAT10a-RFP and LuPAT10b-RFP, which were expressed under the control of a shortened and thereby less active Adh promoter (pAdhΔ). As a control, a further tonoplast associated enzyme from *A. thaliana*, PAT11, which was shown to be active towards a pseudosubstrate (Batistič, 2012), was expressed under the control of the same pAdhΔ promoter and used as the specificity control. Under these conditions, Mn-GFP efficiently targeted to the yeast plasma membrane in presence of either LuPAT10a or LuPAT10b, but not in presence of the AtPAT11 enzyme (Figure 6), further indicating that M is most likely modified by the LuPAT10 enzymes.

**Figure 6:**
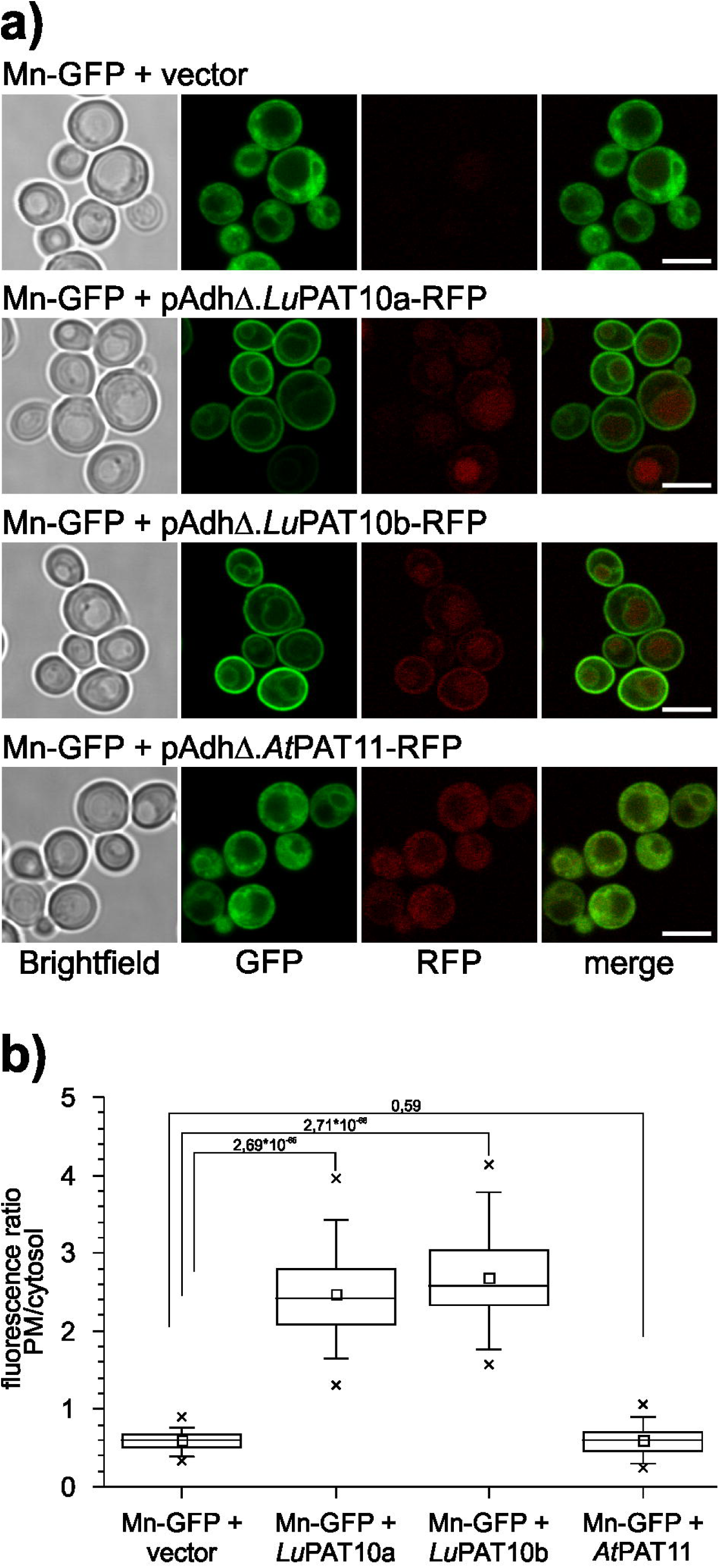
Flax PAT10 specifically mediates the association of Mn with yeast membranes. **a)** Microscopic analysis of yeast cells expressing Mn-GFP + vector (first row), flax PAT10a (LuPAT10a, second row), LuPAT10b (third row) and *A. thaliana* PAT11 (AtPAT11, last row). PATs were expressed under the control of a week pAdhΔ promoter (shortened Adh promoter). Mn efficiently associates with the plasma membrane in yeast cells co-expressing LuPAT10a or LuPAT10b. In presence of AtPAT11-RFP, Mn was not efficiently targeted to the plasma membrane, like in the vector control. Brightfield image is shown in the first picture, GFP in the second, RFP in the third, the fluorescence is merged in the last picture. Bar represents 5 µm. **b)** Membrane binding efficiency of Mn-GFP in yeast cells co-expressing PATs under the control of the weak pAdhD promoter was determined by calculating the ratio of fluorescence intensities at the plasma membrane and in the cytosol (fluorescence intensity PM/cytosol). Co-expression of LuPAT10a (mean ratio 2,47± 0,53SD, n= 75) or LuPAT10b (mean ratio 2,68± 0,59SD, n= 77) strongly increases fluorescence ratio of Mn towards the yeast plasma membrane compared to vector control (0,60±0,12SD, n= 75). The co-expression of AtPAT11 does not significantly changes the ratio (mean ratio 0,61±0,18SD; n= 85 cells). P values from student’s t-test for selected pairs (connected by lines) are indicated above the box plots.

Therefore, to further test if also the full-length M protein is associated with yeast membranes by LuPAT10a or LuPAT10b (here under the control of the regular Adh promoter), the localization of M and of the non-S-acylatable M^C19,20A^ variant were microscopically analysed in yeast cells in absence and presence of the active LuPAT10a/b (Fig. **7**, Fig. S4). In yeast cells, full-length M, which was expressed as an intact fusion protein (Fig. S4a), showed a cytoplasmic localization (Fig. **7a**). In contrast to that, co-expression of either LuPAT10a or LuPAT10b resulted in plasma and vacuolar membrane binding of M (Fig. **7b**). On the other hand, M^C19,20A^ did not localize to membranes neither in presence of LuPAT10a (Fig. **7c**, Fig. S4a) nor in presence of LuPAT10b (Fig. **7c**). Finally, full-length M co-expressed with *A. thaliana* PAT11, did not localize to the yeast plasma membrane nor to the vacuolar membrane (Fig. **7d**), indicating that PAT11 is not active towards M.

**Figure 7:**
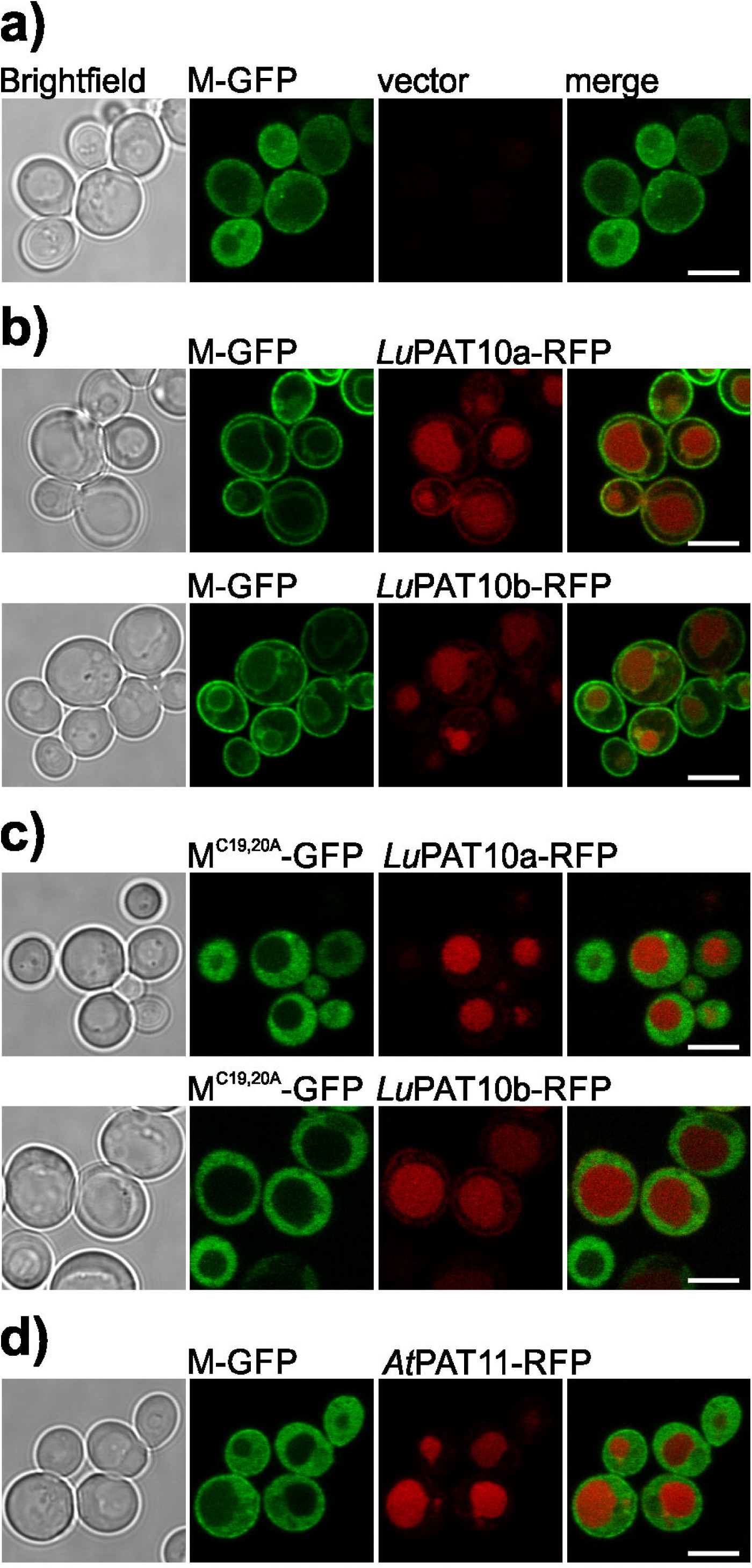
The flax PAT10 enzymes mediate membrane targeting of full-length M in yeast cells. **a)** Full-length M is accumulating in the yeast cytosol (yeast cells co-transformed with empty vector). **b)** M-GFP is associated with the plasma membrane and partially with the vacuolar membrane in yeast cells co-transformed with either LuPAT10a (first row) or LuPAT10b (second row). **c)** Mutated full-length M^C19,20A^-GFP is accumulating in the cytosol of yeast cells co-transformed with either LuPAT10a (first row) or LuPAT10b (second row). **d)** M-GFP is accumulating in the cytosol of yeast cells co-transformed with AtPAT11. All PATs were expressed under the control of the regular pAdh promoter. Brightfield image is shown in the first picture, GFP in the second, RFP in the third, the fluorescence is merged in the last picture. Bar represents 5 µm.

Overall, this shows, that membrane targeting of the M N-terminus and full-length M in yeast cells requires S-acylation of the N-terminal cysteine residues by at least two highly related enzymes, LuPAT10a and LuPAT10b.

### LuPAT10a/b mediates tonoplast targeting of the M N-terminus in the *Arabidopsis* pat10 mutant

As shown above, the N-terminal peptide from M is lipid-modified in yeast cells by flax PAT enzymes, which are homologous to the *Arabidopsis* PAT10. Therefore, to further determine the PAT10 dependent targeting mechanism of Mn in plants, the localization of Mn was analysed in *Arabidopsis* wild-type (WT) and pat10 mutant plants, which were characterized previously (Zhou *et al*., 2013; Qi *et al*., 2013).

First, Mn-GFP was expressed in *Arabidopsis* WT seedlings, together with the plasma membrane marker TM23-RFP (Fig. **8a**). As revealed in *N. benthamiana* plants (Fig. **3b**), Mn-GFP was targeted to the vacuolar membrane, while TM23-RFP associated with the plasma membrane. This demonstrated that the M N-terminus was S-acylated by an *Arabidopsis* PAT, likely mediated by the endogenous PAT10. Subsequently, Mn-GFP and TM23-RFP were co-expressed in homozygous pat10/SALK_018436 mutant seedlings (Qi *et al*., 2013). In contrast to the targeting in WT plant cells, in the pat10 mutant Mn-GFP localized mainly to the plasma membrane as TM23-RFP (Fig. **8b**). Weak fluorescence of Mn was observed on vesicles and at inner membrane structures, indicating that Mn could be still partially bound to the tonoplast in the pat10 mutant. This could be mediated by the activity of other PAT enzymes. Nevertheless, this revealed that PAT10 from *Arabidopsis* is involved in the targeting efficiency of Mn to the vacuolar membrane and that the absence of this enzyme resulted in the relocation of Mn to the plasma membrane.

**Figure 8:**
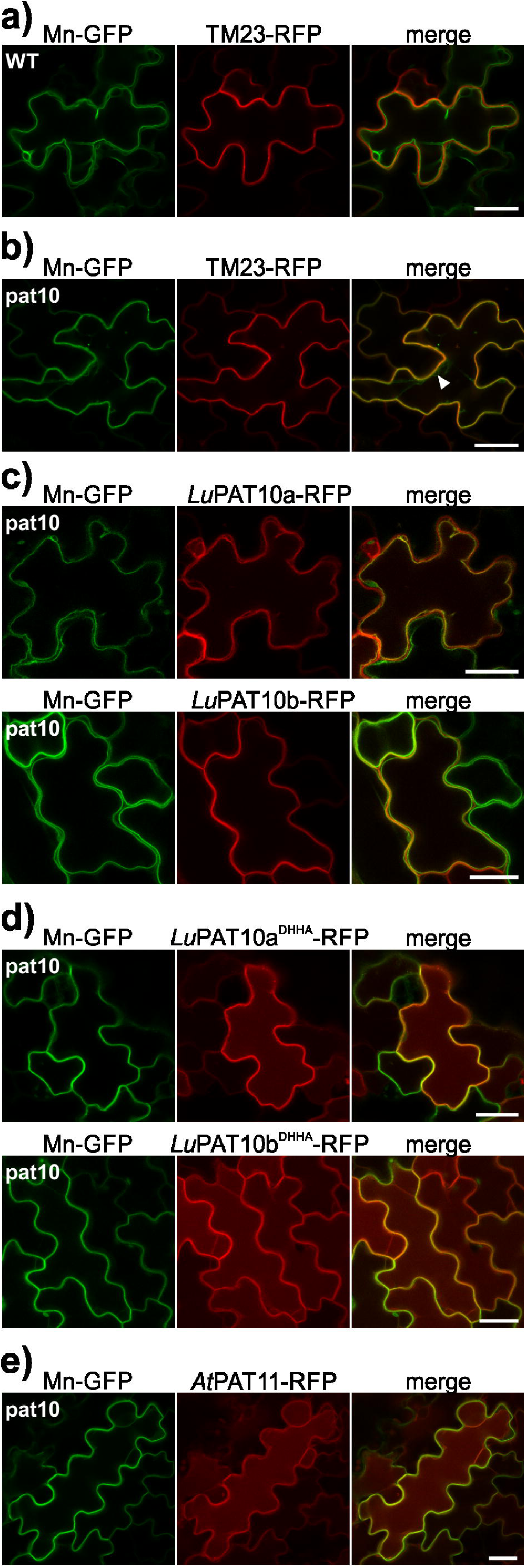
M N-terminus retargets from the plasma membrane to the tonoplast in the *Arabidopsis* pat10 mutant co-expressing active flax PAT10 enzymes. **a)** Mn-GFP and TM23-RFP were transiently expressed in wild-type (WT) *Arabidopsis* seedlings. Mn-GFP displays vacuolar targeting, while the co-expressed TM23-RFP localizes to the plasma membrane **b)** Mn-GFP and TM23-RFP were transiently expressed in pat10 *Arabidopsis* seedlings. Mn-GFP displays plasma membrane targeting, like the co-expressed TM23-RFP. A minor fraction of green fluorescence is still observable at inner membrane structures (arrowhead). **c)** Mn-GFP was transiently co-expressed with LuPAT10a-RFP (first row) and LuPAT10b-RFP (second row) in pat10 *Arabidopsis* seedlings. In the presence of both enzymes, Mn-GFP is targeted to the vacuolar membrane. Both enzymes display a plasma membrane targeting, as well as tonoplast binding. **c)** Mn-GFP was transiently co-expressed with LuPAT10a^DHHA^-RFP (first row) and LuPAT10b^DHHA^-RFP (second row) in pat10 *Arabidopsis* seedlings. Mn-GFP is targeted to the plasma membrane and co-localizes with the inactive enzymes. Cells expressing the inactive enzymes further display a certain red fluorescence within the vacuole. d) Mn-GFP was transiently co-expressed with AtPAT11-RFP in pat10 *Arabidopsis* seedlings. Mn-GFP is targeted to the plasma membrane, while AtPAT11-RFP is associated with the tonoplast and is accumulating within the vacuole. The first image shows GFP, the second image RFP and both fluorescences are merged in the third image. Bar represents 20 µm.

TM23 was then replaced by LuPAT10a or LuPAT10b and co-overexpressed with Mn-GFP in the pat10 mutant plant. As shown, Mn-GFP is efficiently targeted back to the vacuolar membrane in the presence of LuPAT10a or LuPAT10b (Fig. **8c**), showing that these enzymes mediate a similar function as *Arabidopsis* PAT10. Interestingly, these analyses further revealed that LuPAT10a and LuPAT10b mainly targeted to the plasma membrane while only a weak fluorescence was observed at the tonoplast. This is different to *Arabidopsis* PAT10, which is targeted to the vacuolar membrane and Golgi vesicles (Fig. S5a) (Batistič, 2012; Zhou *et al*., 2013; Qi *et al*., 2013). Therefore, to exclude a specific mistargeting of LuPAT10s in *Arabidopsis* cells, LuPAT10a/b-RFP and Mn-GFP were transiently co-overexpressed in cotyledons of flax plants (Fig. S5b). However, this approved that also in flax cells LuPAT10a and LuPAT10b-RFP mainly targets to the plasma membrane, and only weakly associates with the tonoplast membrane. On the other hand, Mn-GFP mainly targets to the vacuolar membrane in flax cells, as observed in *N. benthamiana* and *Arabidopsis* plants. Overall, this rather excludes a specific mistargeting of the tested proteins due to expression in heterologous plant cells. To further show that the targeting of the M N-terminus to the tonoplast is mediated by an active LuPAT10 enzyme, Mn-GFP was co-overexpressed with the inactive LuPAT10a ^DHHA^ or LuPAT10b ^DHHA^ mutant (Fig. **8d**). Moreover, the *Arabidopsis* PAT11-RFP was co-overexpressed to determine the specificity of the PAT10 mediated Mn targeting to the vacuolar membrane (Fig. **8e**). However, in presence of the inactive LuPAT10 enzymes (Fig. **8d**) or active AtPAT11 (Fig. **8e**) Mn-GFP was not targeted to the vacuolar membrane and the fusion protein remained mainly at the plasma membrane. Moreover, similarly to the active LuPAT10 enzymes, LuPAT10a/b mutant protein accumulated at the plasma membrane (Fig. **8d**), while AtPAT11-RFP localized to the tonoplast and further accumulated within the vacuole (Fig. **8e**).

Taken together, these results show that tonoplast targeting of Mn in *Arabidopsis* is mainly facilitated by the endogenous PAT10. The lack of this activity results in plasma membrane relocation of Mn, which is in accordance to observations when using the S-acylation inhibitor 2-BrP (Fig. **3**). Importantly, co-expression of the active flax PAT10 enzymes in the *Arabidopsis* pat10 mutant complemented the tonoplast localization of Mn, although both enzymes mainly targeted to the plasma membrane. This approved that the two LuPAT10 enzymes could be involved in the S-acylation and targeting mechanism of M.

## Discussion

Lipid modifications, like N-myristoylation and S-acylation, are crucial for correct targeting and function of several R proteins. This predominantly mediates the localization to the plant plasma membrane, which naturally represents the first cellular defence barrier against a pathogen (Boyle & Martin, 2015). The R protein variant M from flax represents a particular exception since it is targeted to the vacuolar membrane. This work shows that S-acylation of N-terminal cysteine residues is involved in the efficient targeting of M to the tonoplast. Moreover, an intact N-terminus is required to activate an HR by M in the presence of a pathogen effector, indicating that S-acylation and correct targeting of M is a prerequisite for the accurate R protein function.

The N-terminal targeting signal of M exhibits overlapping structural features which were previously identified in the S-acylated, tonoplast localized CBLs2/3/6 from *Arabidopsis*, but which are apart from that structurally unrelated to M (Takemoto *et al*., 2012; Batistič *et al*., 2012; Zhang *et al*., 2017). Accordingly, mutations of the N-terminal cysteines in M or within the isolated N-terminal S-acylation domain Mn resulted in cytoplasmic accumulation of the GFP fusions, which is comparable to observations using CBL cysteine mutants (Batistič *et al*., 2012; Zhang *et al*., 2017). Similarly to that, the chemical block of S-acylation by the inhibitor 2-BrP also resulted in cytoplasmic accumulation of full-length M, pointing that M membrane binding is mediated by S-acylation. Surprisingly, regardless of whether the short Mn or a longer Mn_30_ fragment was used, the isolated S-acylation domain of M did not accumulate in the cytosol in presence of 2-BrP like full-length M or as it was observed for the isolated CBL2 N-terminus (Batistič *et al*., 2012). In striking contrast, both variants of the M N-terminal domain, which were used here, remained at membranes and in particular, the fragments showed a relocation to the plasma membrane. However, this “translocation efficiency” was not identical for both fragments. While the shorter Mn peptide nearly exhibited full translocation, a large fraction of Mn30 remained at the tonoplast, indicated by the strong fluorescence observed at the inner membrane. At the moment, this difference in the “translocation efficiency” cannot be finally clarified. One the one hand, it is likely that the longer Mn30 fragment is more efficiently S-acylated, and therefore exhibits a weaker effect on the chemical blocking by 2-BrP. Additionally, it could be that the amino acids 24-30 somehow further promotes the binding to the vacuolar membrane, directly or indirectly. Nevertheless, while the N-terminal S-acylation domain of M remained at membranes, full-length M showed a strong cytosolic accumulation in presence of 2-BrP. This could indicate that other regions of the full-length multi-domain protein might influence the function of the N-terminal S-acylation domain. In fact, functional analysis on R proteins revealed extensive intra-domain interactions, and it is further proposed that the N-terminal domains of the R proteins can get in proximity to the C-terminal end which results in an overall closed conformation (Takken & Goverse, 2012). On one hand, this could be crucial to keep the protein in a resting state (Takken & Goverse, 2012; El Kasmi *et al*., 2017), but this intra-molecular interaction might also influence the membrane-binding potential of non-S-acylated N-terminal M domain. Nevertheless, these analyses revealed that the N-terminus of M could represent a membrane-binding domain even in absence of a lipid modification event and thereby likely directs S-acylation of M. In fact, S-acylation requires particular preconditions to occur, like hydrophobic domains or other lipid modifications (Turnbull & Hemsley, 2017). As in tonoplast-localized CBLs2/3/6, the N-terminus of M lacks an N-myristoylation signal, and it is still unclear how the S-acylation of these proteins is initialized. Here, the autonomous membrane binding of the N-terminus could represent the initial step in the targeting and S-acylation of M but also of the related N-terminal domains found in CBLs. This initial membrane binding depends on the presence of intact cysteine residues, which could rely on the certain hydrophobic nature of these amino acids (Nagano *et al*., 1999). However, while the isolated N-terminus of M alone allows autonomous membrane binding when fused to a rather small protein like GFP, this domain appears to be less efficient in membrane binding within the bulkier full-length M protein and therefore requires additional S-acylation to convey stable membrane localization. If this targeting involves lipid modification of both N-terminal cysteines remains open, but the results presented revealed that lipid modification of cysteine 19 is sufficient to mediate tonoplast binding, at least then using the short Mn S-acylation domain. This does not exclude that also cysteine 20 is further S-acylated, but it could be that the latter cysteine is rather involved in promoting the reactivity of cysteine 19 and thereby is increasing the overall S-acylation efficiency (Parente *et al*., 1985) and thereby could be important for the effective lipid modification especially of the full-length M protein. However, at the moment, it cannot be fully excluded that also other domains of M might contribute to membrane binding and to the targeting efficiency, especially to the vacuolar membrane.

Finally, the sequence and targeting similarities between M and CBL2/3/6 N-termini further implicated that the lipid modification of these proteins involves related enzymes. In *A. thaliana*, CBLs2/3/6 localization to the vacuolar membrane requires the function of the tonoplast/Golgi bound PAT10 enzyme, which to date was not observed at the plasma membrane (Batistič, 2012; Zhou *et al*., 2013; Qi *et al*., 2013). However, as observed after chemical blocking of S-acylation, the Mn peptide was targeted to the plasma membrane in the pat10 mutant, showing that this enzyme is involved in the correct localization of Mn. This indicated that in flax a similar mechanism is involved in the lipid modification of M. In accordance to that, two highly related flax PAT10 enzymes, homologous to *A. thaliana* PAT10, were identified in the flax genome and can complement the mislocalization of Mn in the pat10 mutant. This demonstrates that these enzymes could mediate S-acylation of M. Moreover, both enzymes thioesterified Mn and mediated plasma membrane binding in yeast cells. Importantly, also full-length M was efficiently targeted to membranes in yeast cells in the presence of the flax PAT10s but not by the *Arabidopsis* PAT11, showing that this localization requires active and specific enzymes. Overall, this indicates that these enzymes are indeed involved in the lipid modification of M, but it cannot fully be excluded that also other enzymes are mediating M S-acylation and targeting. Surprisingly, in contrast to *Arabidopsis* PAT10, the flax PAT10a and PAT10b enzymes were mainly observed at the plasma membrane and displayed an only partial association with the vacuolar membrane. Interestingly, the plasma membrane localization coincides with the “mistargeting” of the Mn peptide in the pat10 mutant, or in presence of 2-BrP. This could indicate that the flax PAT10 enzymes modify proteins at various membranes, but especially at the plasma membrane. These results indicate that besides a direct trapping of proteins to the vacuolar membrane by S-acylation as previously proposed for CBL proteins (Batistic, 2012), a further pathway could exist in flax cells and potentially also in other plant organisms, where proteins are lipid-modified at the plasma membrane and then transported to the vacuolar membrane.

Overall, this work shows that the N-terminus of M represents an S-acylation domain and that this modification is likely required for M targeting and function at the tonoplast. This modification could be mediated by a class of enzyme which is known to be involved in the lipid modification of *A. thaliana* tonoplast targeted CBLs. These two flax PAT10 enzymes seem to be mainly targeted to the plasma membrane and partially to the vacuolar membrane. The results presented here indicate that in flax certain proteins could be lipid-modified at the plasma membrane and subsequently are transported to the tonoplast. Therefore, in order to further determine the role and interplay of M S-acylation and targeting during pathogen response, it would be of particular interest to investigate the S-acylation of M and the localization of the flax PAT10 enzymes in the presence of pathogens or pathogen effectors in the future.

## Acknowledgements

Dr. Bauxiu Qi (University of Bath, England) kindly provided pat10 mutant seeds. This work was supported by the Deutsche Forschungsgemeinschaft (DFG BA4742/1-1 and 1-2).

## Supporting information

### Supplementary figures

Figure Sl: Targeting of TM23-RFP in presence of 2-BrP

Figure S2a: Targeting of Mn_30_-GFP in absence and presence of 2-BrP

Figure S2b: S-acylation of Mn_30_-GFP

Figure S3a: Phylogenetic tree of flax and *Arabidopsis* PAT proteins

Figure S3b: Alignment of flax and *Arabidopsis* PAT10

Figure S3c: Similarity and identity between LuPAT10a/b and AtPAT10

Figure S4a: Protein integrity of full-length M-GFP and M^C19,20A^-GFP expressed in yeast cells

Figure S4b: Localization of full-length M to the yeast plasma membrane

Figure S5a: Microscopic analyses of *Arabidopsis* PAT10 and flax PAT10a/b in *A. thaliana* seedlings

Figure S5b: Localization of flax M-n and PAT10a/b in flax seedlings

### Supplementary Material and Method

Table S1: Primers used in this work

Construct generation

Supplementary References

