## Supplemental Figures for "S-acylation is involved in tonoplast targeting of flax resistance protein M"

#### Supporting Information

#### Figure S1

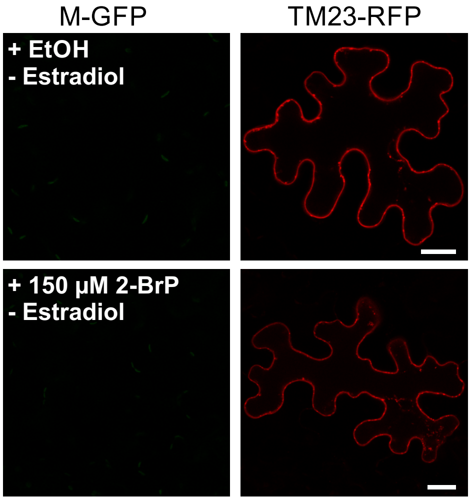


Targeting of TM23-RFP in presence of 2-BrP

TM23-RFP and M-GFP were co-transformed in *N. benthamiana* leaves. M-GFP (shown in the left images), under the control of the Estradiol promoter, was not expressed since Estradiol was omitted. TM23-RFP (shown in the right images) is targeted to the plasma membrane and vesicles in absence of 2-BrP (EtOH solvent control in the upper images) and presence of 150 µM 2-BrP (lower images). Bar represents 20 µm.

### Supporting Information

#### Figure S2

a)

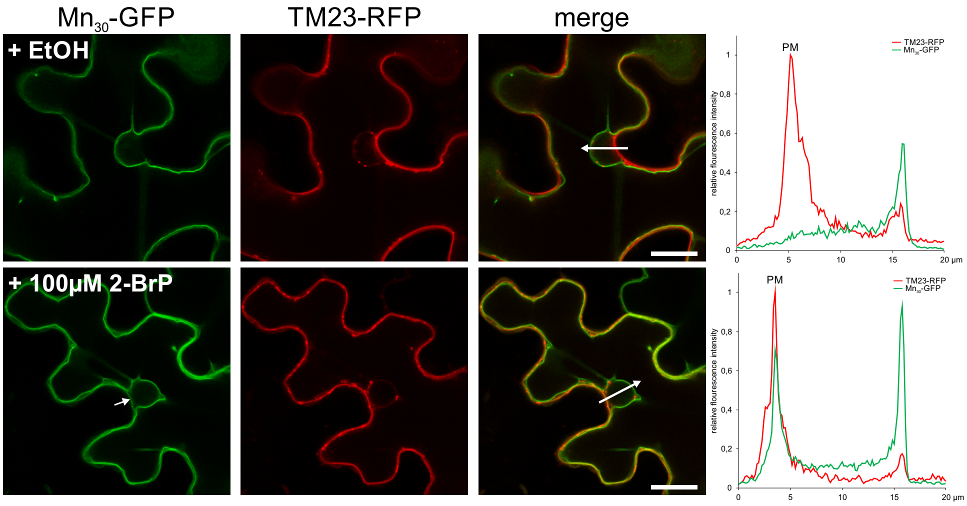


a) Targeting of $Mn_{30}$-GFP in absence and presence of 2-BrP.

The first 30 amino acids of M fused to GFP ($Mn_{30}$-GFP) (first column), was co-expressed with the plasma membrane marker TM23-RFP in *N. benthamiana* leaves. Samples were either incubated in the presence of EtOH control solution (first row) or 100 µM 2-BrP (second row). In presence of the EtOH control solution, Mn30-GFP associates with the tonoplast and not with the plasma membrane, but partially targets to the plasma membrane in presence of 2-BrP (indicated by an arrow in the $Mn_{30}$-GFP image). The fluorescence intensity was measured within the region of interest (indicated by the arrow in the merged picture) and is depicted in the right graph (PM indicates the position of the plasma membrane). The scale bars depicted in the overlay images are 20 µm.

b)

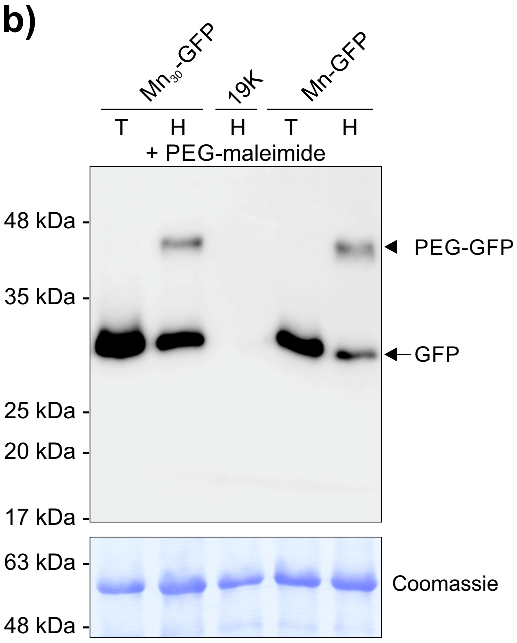


b) S-acylation of $Mn_{30}$-GFP

S-acylation of $Mn_{30}$-GFP (samples left) was determined by an S-acyl-PEG switch assay and compared to Mn-GFP (samples right). $Mn_{30}$-GFP and Mn-GFP were expressed in *N. benthamiana* leaves. Proteins were extracted in the presence of NEM (block of free cysteine residues) and subsequently incubated in either Tris (T) or Hydroxylamine solution (H) in the presence of PEG-maleimide. A leaf expressing only the helper plasmid 19K (incubated with Hydroxylamine) was used as an antibody specificity control (sample in the middle). Recombinant GFP proteins were detected in all samples (except the 19K control) at the expected size (~30 kDa; upper immunoblot). In both $Mn_{30}$-GFP and Mn-GFP samples incubated with Hydroxylamine, a second shifted protein is present indicating S-acylation of both proteins. Coomassie staining of the rbcL protein (lower blot) is shown as a loading control. The position of a protein size marker is indicated.

#### Supporting Information

#### Figure S3

**a)**

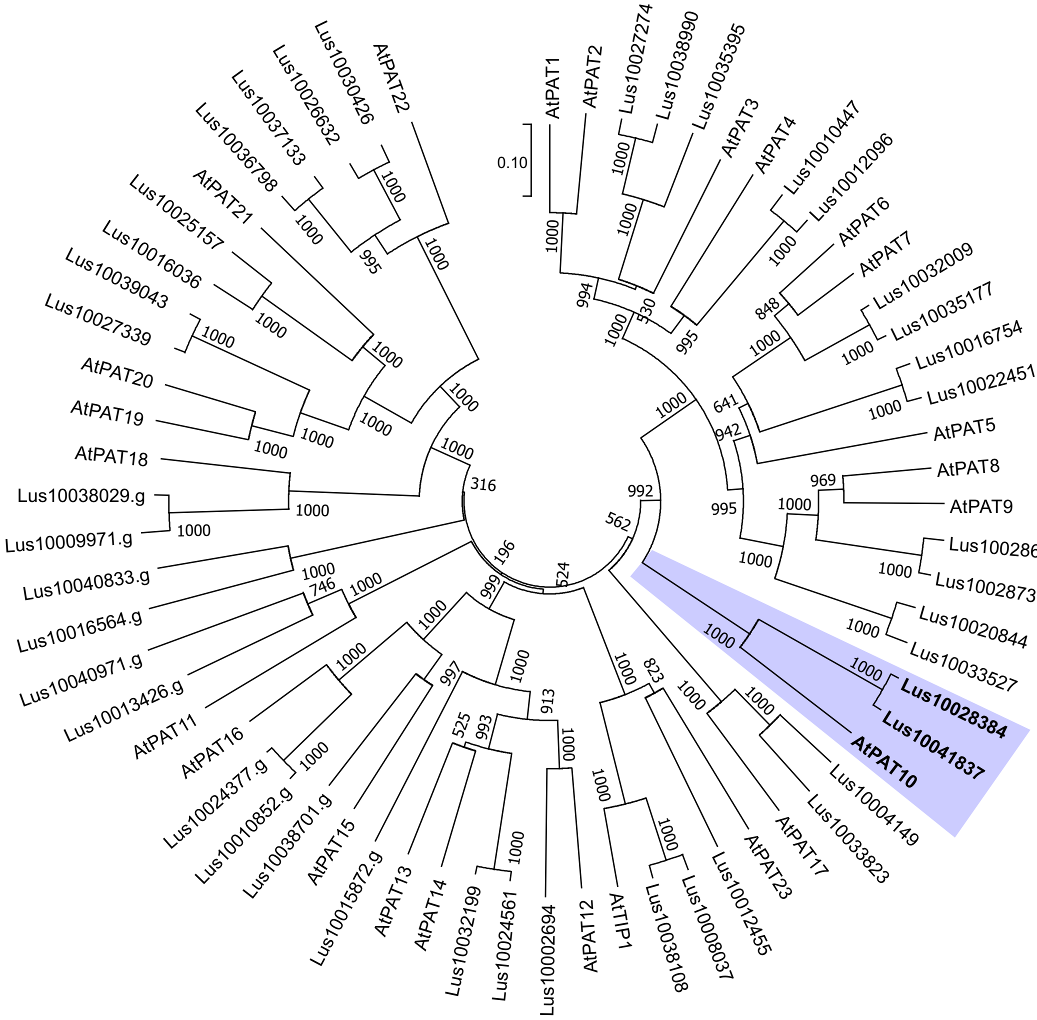


a) Phylogenetic tree of flax and *Arabidopsis* PAT proteins

Flax PAT protein sequences were obtained from Phytozome and aligned against *Arabidopsis* PATs using ClustalX. A radial, phylogenetic tree was constructed by a neighbour-joining method. Two flax sequence annotations (Lus10028384 and Lus10041837) branched together with *Arabidopsis* PAT10 (*At*PAT10) (highlighted by the purple background). All other sequences branched with other *Arabidopsis* PATs. Bootstrap factors are given at the nodes.

#### Supporting Information

#### Figure S3 (continued)

**b)**

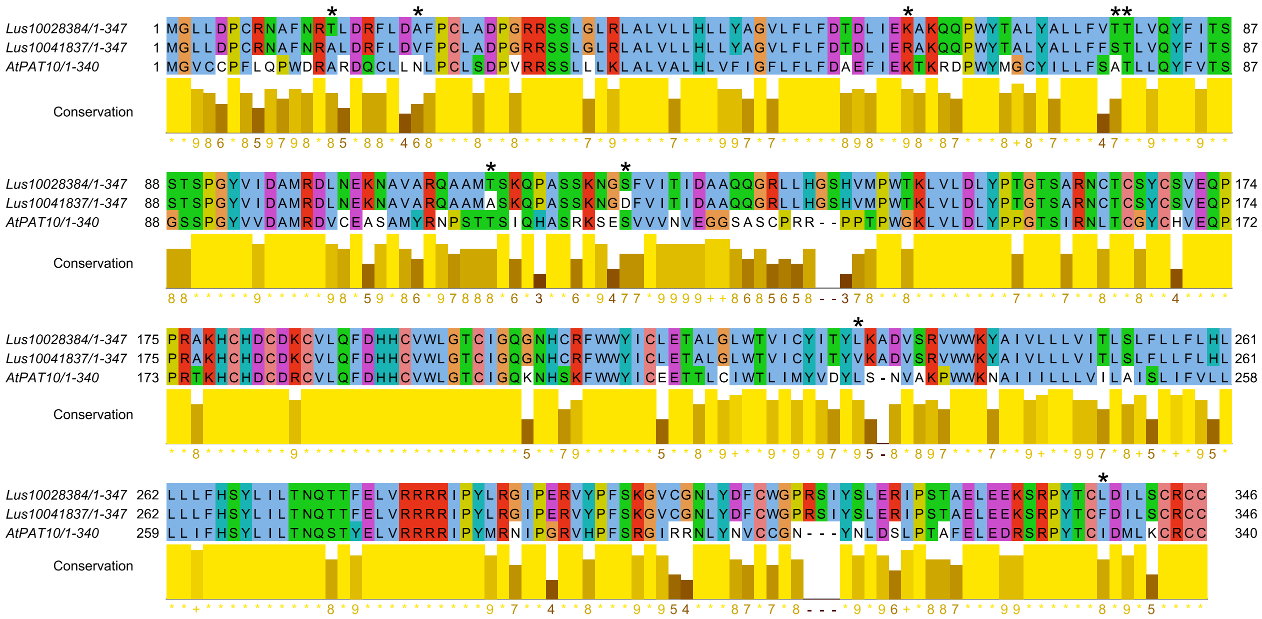


b) Alignment of flax and *Arabidopsis* PAT10

Alignment of the two *L. ussitatissimum* PAT10 annotations (*Lu*PAT10a/Lus10028384 and *Lu*PAT10b/Lus10041837) and *A. thaliana* PAT10 (*At*PAT10). The amino acid discrepancies between *Lu*PAT10a and *Lu*PAT10b are indicated by an asterisk. Amino acids are coloured using the Clustal-X colour scheme. Yellow columns below the sequences indicate amino acid conservation.

**c)**

| % similarity \ % identity | *Lu*PAT10a | *Lu*PAT10b | *At*PAT10 |
|---|---|---|---|
| *Lu*PAT10a | - | 97,4 | 58,1 |
| *Lu*PAT10b | 98,3 | - | 57,2 |
| *At*PAT10 | 73,7 | 73,4 | - |

c) Similarity and identity between *Lu*PAT10a/b and *At*PAT10

Protein similarity (orange background) and identity (green background) compared between *Lu*PAT10a, *Lu*PAT10b and *At*PAT10.

#### Supporting Information

#### Figure S4

**a)**

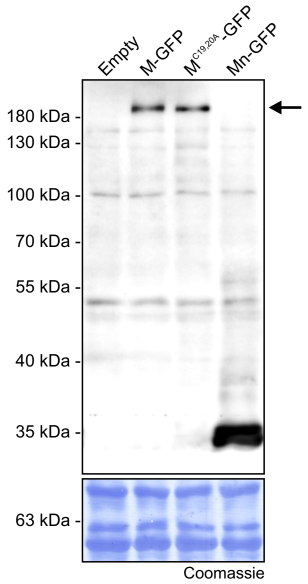


a) Protein integrity of full-length M-GFP and $M^{C19,20A}$-GFP expressed in yeast cells

Full-length M and $M^{C19,20A}$, fused to GFP (M-GFP and $M^{C19,20A}$-GFP, respectively), were expressed in yeast cells and the protein integrity was determined by western blot analysis. M-GFP, as well as $M^{C19,20A}$-GFP (lane 2 and 3), appeared at a size about ~180 kDa (arrow) compared to the size marker used (indicated on the left, calculated size of M-GFP= 175 kDa), showing that both proteins are expressed as intact full-length proteins. Other bands which appeared at smaller sizes, were also visible in protein extracts from yeast cells transformed with vector only (first lane), as well as from yeast cells transformed with Mn-GFP (which appeared at ~35 kDa, last lane), indicating that these additional proteins bands unspecifically cross-reacted with the antibodies used. As a loading control, the western was stained with Coomassie blue (shown below).

**b)**

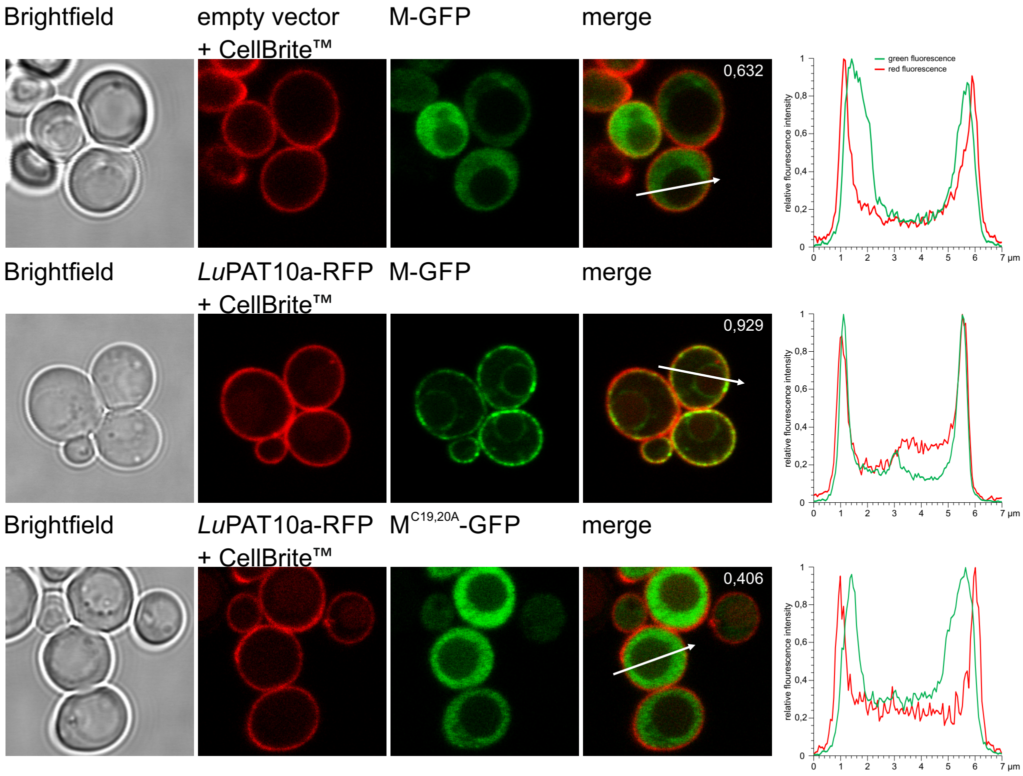


b) Localization of full-length M to the yeast plasma membrane

Plasma membrane localization of full-length M to the yeast plasma membrane was further quantified. To clearly determine the position of the plasma membrane, yeast cells transformed with M-GFP and vector control (first row), M-GFP and LuPAT10a-RFP (second row), and $M^{C19,20A}$-GFP co-transformed with *Lu*PAT10a-RFP (third row) were stained with CellBrite™ Fix 555 for at least 10 min. The first image presents the brightfield picture, the red fluorescence from the CellBrite™ stain is shown in the second image. Please note that cells which co-expressed *Lu*PAT10a could show some, but weaker background fluorescence from the RFP fusion. GFP is shown in the third image, an overlay is shown in the last image. The fluorescence intensity was measured within the region of interest (indicated by the arrow in the merged picture) and is depicted in the graph on the right (length of ROI is 7 µm). In the presence of *Lu*PAT10a, M co-localizes with the CellBrite™ stained plasma membrane, indicated by the overlay of the red and green fluorescence peaks. In contrast, the maxima of the green fluorescence are shifted compared to the red fluorescence peaks in M + vector as well as in $M^{C19,20A}$-GFP + LuPAT10a-RFP. The number within the merged image represents the Pearson correlation coefficient of the two fluorescences within the ROI.

#### Supporting Information

#### Figure S5

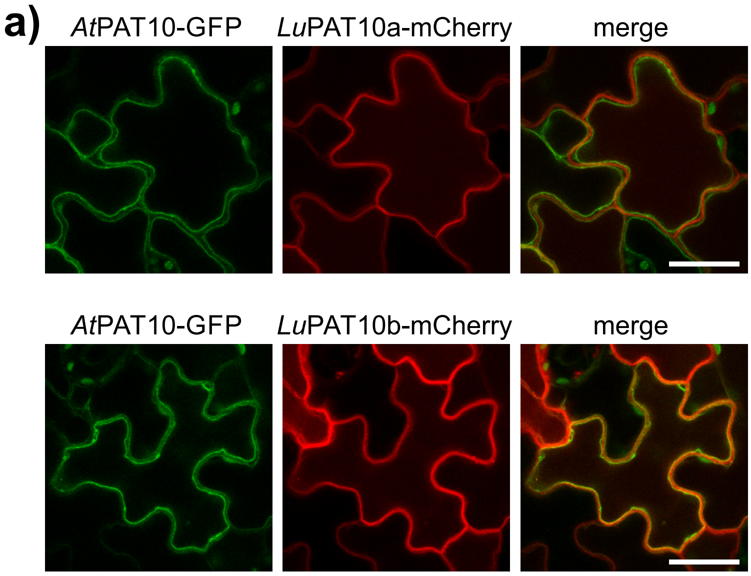


a) Localization of *Arabidopsis* PAT10 and flax PAT10a/b in *A. thaliana* seedlings

*A. thaliana* PAT10 fused to GFP (*At*PAT10-GFP) and *Lu*PAT10a, as well as *Lu*PAT10b, fused to RFP (*Lu*PAT10a/b-RFP), were transiently co-expressed in *Arabidopsis* WT seedlings. *At*PAT10 (first image) is targeted to the tonoplast and vesicles, while *Lu*PAT10a (second image, first row) and *Lu*PAT10b (second row) mainly target to the plasma membrane. A merge of the fluorescences is shown in the third image. The scale bar in the merged picture represents 20 µm.

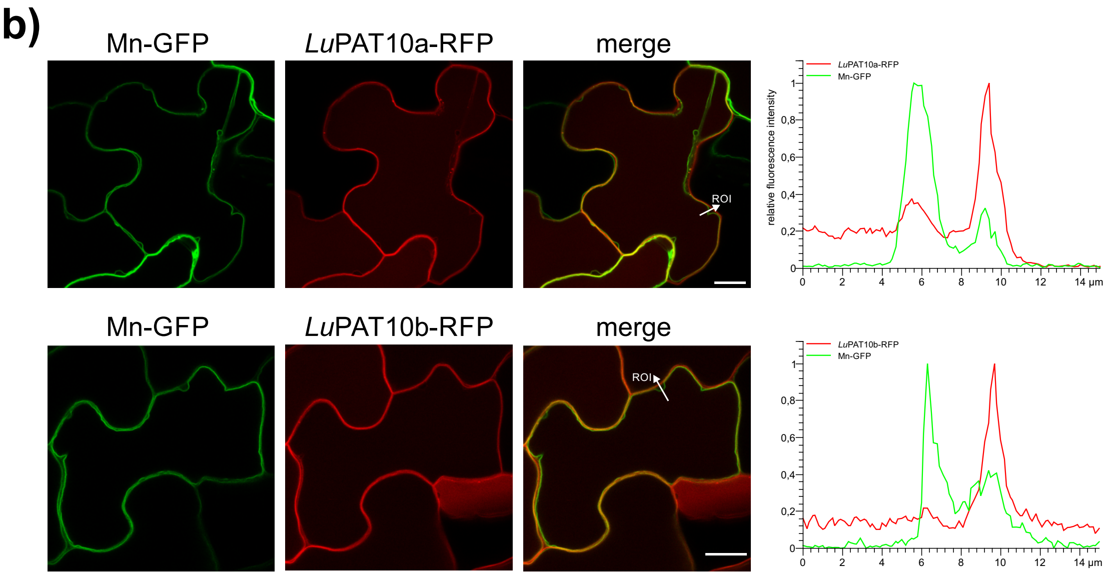


b) Localization of flax M-n and PAT10a/b in flax seedlings

Flax M N-terminus fused to GFP (Mn-GFP) and flax PAT10a/b fused to RFP (*Lu*PAT10a/b-RFP) were transiently co-expressed in flax seedlings. Mn-GFP (first image) is mainly targeted to the vacuolar membrane, while *Lu*PAT10a (second image, first row) and *Lu*PAT10b (second image, second row) mainly targeted to the plasma membrane and partially to the vacuolar membrane. Fluorescence intensity was determined within the indicated region of interest (arrow) and is depicted in the right graph. The scale bars in the merged images represent 20 µm.
