## Supplemental Material for "S-acylation is involved in tonoplast targeting of flax resistance protein M"

### Supporting Information

### Supplementary Material and Method

**Table S1**: Primers used in this work

| **Name** | **Sequence** |
|---|---|
| MnGFPforXbaI | aaaTCTAGAtgaaaaatgtcttatcttagagatgttgctactgctgttgctcttcttcttgataatctttgttgtggaagacctagtaaaggagaagaacttttcac |
| MnCxxAGFPforXbaI | aaaTCTAGAtgaaaaatgtcttatcttagagatgttgctactgctgttgctcttcttcttgataatcttgctgctggaagacctagtaaaggagaagaacttttcac |
| MnC19AGFPforXbaI | aaaTCTAGAtgaaaaatgtcttatcttagagatgttgctactgctgttgctcttcttcttgataatcttgcttgtggaagacctagtaaaggagaagaacttttcac |
| MnC20AGFPforXbaI | aaaTCTAGAtgaaaaatgtcttatcttagagatgttgctactgctgttgctcttcttcttgataatctttgtgctggaagacctagtaaaggagaagaacttttcac |
| Mn30aaGFPforXbaI | aTCTAGAatgagttatttgagagacgttgctactgcggttgccttgcttcttgacaatttgtgctgtgggagaccaaatctcaacaatgacaacgagagtaaagg agaagaacttttcac |
| GFPrevSacI | tttGAGCTCttacttgtacagctcgtccatg |
| LuPAT10forXbaI | ttttTCTAGAtgaaaaatggggcttctcgatccatgc |
| LuPAT10revXmaI | aaaaCCCGGGgcaacagcggcagctcaaaatat |
| LuPAT10DHHAfor | gtgttctccagtttgatcatcacGCCgtttggcttgggacatgcattg |
| LuPAT10DHHArev | caatgcatgtcccaagccaaacGGCgtgatgatcaaactggagaacac |
| AtPAT11for | tttTCTAGAaaaatggaagattcttcccagggg |
| AtPAT11rev | aaaaCCCGGGtgtcttatgtctcttcctcaag |
| MfullForXhoI | ttttCTCGAGtgaaaaatgagttatttgagagacgttgc |
| MpartIRev | ctaatggaaattcaaactggacctcctgagctccttctgttttcaagactttg |
| MpartIIFor | caaagtcttgaaaacagaaggagctcaggaggtccagtttgaatttccattag |
| MfullRevXmaI | ttttCCCGGGcttatatttctcgtccggcc |
| MfullCxxAForXhoI | ttttCTCGAGtgaaaaatgtcttatcttagagatgttgctactgctgttgctcttcttcttgataatcttgctgctgggagaccaaatctcaacaatg |
| AvrMdspforXbaI | ttttTCTAGAtgaaaaatgcaccccatgaactcagcaaaac |
| AvrMArevXmaI | aaaaCCCGGGcatgtctggagatttcaatatcttg |
| AvrMrevXmaI | aaaaCCCGGGattgttttcttgataagacagggttc |

Restriction sites are underlined and in capital letters; mutagenesis sites are given in capital letters

### Construct generation

M-n:GFP and M-nC19,20A were amplified with forward primers MnGFPfor/MnGFPCxxAfor and reverse primer GFPrevSac, respectively, using GFP cDNA as a template (Batistič *et al.*, 2008). Accordingly, M-n:The PCR products were inserted in pGPTVII plasmids (Walter *et al.*, 2004) and expression is driven using an UBI10 promoter and Hsp18.2 terminator (Grefen *et al.*, 2010; Nagaya *et al.*, 2010). cDNA clones for Lus10028384/LuPAT10a as well as for Lus10041837/LuPAT10b were generated by PCR, using the generated cDNA mix as template and using the primer-pair LuPAT10forXbaI-LuPAT10revXmaI. The amplificates were integrated into a pDE1002 vector and the specific cDNAs for Lus10028384 and Lus10041837 were isolated from several *E. coli* clones and verified by sequencing. LuPAT10a/b$^{DHHA}$ (cysteine 194 to alanine mutation) were generated by site directed mutagenesis (Higuchi *et al.*, 1988), using primers LuPAT10DHHAfor and LuPAT10DHHArev and the respective LuPAT10 cDNAs as template. WT LuPAT10a/b and the mutagenized LuPAT10a/b$^{DHHA}$ were inserted into pDE1002 plasmid and fused with the red fluorescence protein (RFP) mCherry. Expression is driven by an UBI10 promoter and Hsp18.2 terminator. PAT11 was amplified from a cDNA as described (Batistič, 2012), inserted into the pDE1002 plasmid and fused with the mCherry RFP. Expression is driven by an UBI10 promoter and Hsp18.2 terminator. For expression in yeast cells, M-n:GFP and M-nC19,20A:GFP were inserted in pVT-U plasmid (Vernet *et* *al.*, 1987). Expression is driven by the Adh promoter and Adh terminator. LuPAT10 and LuPAT10DHHA were fused with mCherry and inserted into pGVac8Leu plasmid, a modified pGAD.GH vector (Batistič, 2012). Expression is driven using the Vac8 promoter (Hou *et al.*, 2009) and Adh terminator. Additionally, LuPAT10 was expressed using the Adh promoter or using a fragment of the Adh promoter (pAdhΔ, nucleotides -410 - -1). For AtPAT11 in yeast cells, the PAT11 reading frame (Batistič, 2012) was inserted into the modified pGAD.GH vector, fused with mCherry and expression was driven by the Adh promoter/terminator. All newly generated plasmids were verified by sequencing.

For generating full-length flax M, two DNA fragments were synthesized at BioCat GmbH (Heidelberg, Germany), based on the published sequence (GenBank: U73916.1) (Anderson, 1997). The fragments cover the nucleotides 1-2422 and 2423-3963 of the ORF, respectively. The fragments were amplified by PCR using primers MfullForXhoI-MpartIRev (fragment 1) and primers MpartIIFor-MfullRevXmaI (fragment 2), respectively. The particular PCR products were pooled and used as templates in a second PCR (using forward MfullForXhoI and reverse MfullRevXmaI primer), to fuse the fragments to a complete ORF. This PCR product was inserted into a pGPTVII vector containing the XVE inducible expression cassette (Schlücking *et al.*, 2013), thereby fused to GFP N-terminally. Similar to that, full-length MC19,20A mutant was generated by PCR (amplified with MfullCxxAForXhoI-MfullRevXmaI, using full length M as template) and inserted into the inducible pGPTVII plasmid.

AvrM-AΔSP and avrMΔSP, which lack the proposed signal peptide (ΔSP) as described (Catanzariti *et al.*, 2006), were amplified from synthetic DNA templates using primers AvrMdspforXbaI-AvrMArevXmaI/AvrMrevXmaI, respectively. The synthetic templates were generated at BioCat GmbH (Heidelberg, Germany), based on the published nucleotide sequences (AvrM-A GenBank: DQ279864; avrM GenBank: DQ279870). The PCR amplificates were inserted into the pDE1002 plasmid, thereby fused to the N-terminus of the RFP mCherry, and expression is driven by the UBI10 promoter and Hsp18.2 terminator.

Following constructs were generated previously: GFP (Batistič *et al.*, 2008), TM23:RFP (Batistič *et al.*, 2012), TPK1:RFP and PAT10:GFP (Batistič, 2012).
